# Calcium-dependent regulation of neuronal excitability is rescued in Fragile X Syndrome by a tat-conjugated N-terminal fragment of FMRP

**DOI:** 10.1101/2024.03.01.583006

**Authors:** Xiaoqin Zhan, Hadhimulya Asmara, Paul Pfaffinger, Ray W. Turner

## Abstract

Fragile X Syndrome arises from the loss of Fragile X Messenger Ribonucleoprotein (FMRP) needed for normal neuronal excitability and circuit functions. Recent work revealed that FMRP contributes to mossy fiber LTP by adjusting Kv4 A-type current availability through interactions with a Cav3-Kv4 ion channel complex, yet the mechanism has not yet been defined. In this study using wild-type and *Fmr1* knockout (KO) tsA-201 cells and cerebellar sections from *Fmr1* KO mice, we show that FMRP associates with all subunits of the Cav3.1-Kv4.3-KChIP3 complex, and is critical to enabling calcium-dependent shifts in Kv4.3 inactivation to modulate A-type current. Specifically, upon depolarization Cav3 calcium influx activates dual specific phosphatase 1/6 (DUSP1/6) to deactivate ERK1/2 (ERK) and lower phosphorylation of Kv4.3, a signalling pathway that does not function in *Fmr1* KO cells. In *Fmr1* KO mouse tissue slices cerebellar granule cells exhibit a hyperexcitable response to membrane depolarizations. Either incubating *Fmr1* KO cells or *in vivo* administration of a tat-conjugated FMRP N-terminus fragment (FMRP-N-tat) rescued Cav3-Kv4 function and granule cell excitability, with a decrease in the level of DUSP6. Together these data reveal a Cav3-activated DUSP signalling pathway critical to the function of a FMRP-Cav3-Kv4 complex that is misregulated in *Fmr1* KO conditions. Moreover, FMRP-N-tat restores function of this complex to rescue calcium-dependent control of neuronal excitability as a potential therapeutic approach to alleviating the symptoms of Fragile X Syndrome.

**Significance Statement:** Changes in neuronal excitability and ion channel functions have been a focus in studies of Fragile X Syndrome. Previous work identified ion channels that are regulated by FMRP through either protein translation or direct protein-protein interactions. The current study reveals FMRP as a constitutive member of a Cav3-Kv4 complex that is required for a Cav3-DUSP-ERK signalling pathway to increase A-type current and reduce cerebellar granule cell excitability. In *Fmr1* KO cells, Cav3-Kv4 function and calcium-dependent modulation of A-type current is lost, leading to a hyperexcitable state of cerebellar granule cells. Pretreating with FMRP-N-tat restores all Cav3-Kv4 function and granule cell excitability, providing support for FMRP-tat peptide treatment as a potential therapeutic strategy for Fragile X Syndrome.

## Introduction

Fragile X Syndrome (FXS) arises through a CGG repeat expansion in the *Fmr1* gene that blocks expression of Fragile X Messenger Ribonucleoprotein (FMRP), representing the leading monogenic cause of autism spectrum disorders (ASD). Cell and circuit function in FXS is disrupted because FMRP exerts vast influence over protein translation and modulation of ion channels that control neuronal excitability (Contractor et al., 2015; Deng and Klyachko, 2021). The cerebellum acts as an important sensory-motor interface that impacts the activity of cortical systems that contribute to ASD (D’Mello and Stoodley, 2015; Gibson et al., 2023; Gallagher and Hallahan, 2012; Liu et al., 2021). A prominent layer of granule cells receive a massive array of mossy fibre input carrying sensory information that is processed to enable time coding, sensori-motor integration, and predictive expectation (D’Angelo and De Zeeuw, 2009; Garrido et al., 2013; Giovannucci et al., 2017; Sgritta et al., 2017; Wagner et al., 2017). Control over the excitability of granule cells could then exert significant influence on higher levels of circuit function relevant to ASD and FXS.

Membrane excitability in granule cells is known to be highly sensitive to the availability of Kv4 potassium channel mediated “A-type” current (I_A_). Bursts of mossy fibre input to granule cells was shown to reduce I_A_ through a mechanism that involves ERK1/2 (ERK), a kinase increasingly recognized for its role in FXS and ASD (Curia et al., 2013; Osterweil et al., 2013; Asiminas et al., 2019; Murari et al., 2023). A resulting increase in the rate of spike firing then contributes to long-term potentiation of mossy fibre input (Zhan et al., 2020). Cerebellar granule cells also express a Cav3-Kv4 complex that invokes calcium-dependent modulation of Kv4 availability through association with a calcium-sensing KChIP3 protein (Anderson et al., 2010a; Heath et al., 2014; Vierra and Trimmer, 2022). Recently FMRP was found to be closely associated with the Cav3-Kv4 complex (Zhan et al., 2020), yet the manner in which FMRP integrates into the complex, and how ERK is involved in controlling I_A_ and membrane excitability is unknown.

The current study reveals a novel signalling pathway whereby Cav3.1 calcium influx activates the phosphatase DUSP1/6 to reduce activated ERK and phosphorylation of subunits in the complex that increases I_A_ availability. FMRP proves to be critical to calcium-dependent modulation of Kv4.3 as well as the expression levels of DUSP6, where loss of FMRP in *Fmr1* KO mice leads to hyperexcitability in cerebellar granule cells. Yet all aspects of Cav3-Kv4 complex function and granule cell excitability are rescued in *Fmr1* KO cells within 24 hr of reintroducing a tat-conjugated N-terminal fragment of FMRP. The results thus identify a new signalling pathway that depends on FMRP to control the influence of IA on intrinsic excitability of a key cell type in receipt of sensory input.

## Material and Methods

### Cell lines

tsA-201 cells originally derived from a female were purchased from Sigma-Aldrich (Cat# 96121229) and maintained in DMEM supplemented with 10% heat inactivated foetal bovine serum and 1% pen– streptomycin at 37 °C (5% CO2). The calcium phosphate method was used to transiently transfect cDNAs of heterologous proteins. Cells were washed with fresh medium 16 -18 hr after transfection and then transferred to 32 °C (5% CO2) and cultured for up to 36 hr prior to tests. Previous work has determined that tsA-201 cells we use do not endogenously express Cav3.1 or Kv4 channels (X. Zhan, unpublished observations), and have no expression of KChIP3 (**Figure 1-1**). For different experiments, cells were transfected with different combinations of human cDNA Cav3.1 (2.2 µg), Kv4.3 (1.5 µg), KChIP3 cDNA (1.5 µg) as indicated in results and figures. In addition, cells were transfected with eGFP cDNA (1 μg) and human Kir2.1 cDNA (1 μg) for electrophysiology and high potassium stimulation experiments, respectively. CRISPR-Cas9 was used to create an *Fmr1* knockout tsA-201 cell line (**Figure 1-2**). For this three short guide oligonucleotides were synthesised, annealed and cloned into the pSpCas9(BB)-2A-GFP (pX458) vector (Addgene), and constructs were then transfected into tsA201 cells with Lipofectamine 2000 (Invitrogen). After transfection, GFP containing cells were sorted into 96 well plates by flow cytometry. Western blotting and DNA sequencing were used to confirm functional knockout of the *Fmr1* gene. All *Fmr1* knockout tsA-201 cells used in the experiments were maintained from clone 7. To verify that knockout of *Fmr1* did not alter subunit expression, we conducted Western blot on cells coexpressing Cav3.1-Kv4.3-KChIP3 and found no significant difference for any of the subunits between WT and KO cells (**Figure 1-3**).

### Mouse lines

Wild-type (Jackson Lab stock #004828, FVB.129P2-Pde6b+ Tyrc-ch/ AntJ, https://www.jax.org/strain/004828) and *Fmr1* knockout (Jackson Lab stock #004624, FVB.129P2-Pde6b+ Tyrc-ch *Fmr1*tm1Cgr/J, https://www.jax.org/strain/004624) mice were maintained in an Animal Resource Center of the University of Calgary in accordance with guidelines of the Canadian Council of Animal Care. Animals had free access to water and food with a temperature of 20 - 23 °C on a 12 hr light and dark cycle.

### Cerebellar slice preparation

Sagittal cerebellar sections of 250 µm thickness were prepared from P21-P25 male mice as previously described ^19^. Briefly, after isoflurane inhalation anaesthesia animals were decapitated and the cerebella were dissected out and placed in ice-cold artificial cerebrospinal fluid (aCSF) composed of (in mM): 125 NaCl, 25 NaHCO_3_, 25 D-glucose, 3.25 KCl, 1.5 CaCl_2_, 1.5 MgCl_2_ bubbled with carbogen (95% O_2_ and 5% CO_2_) gas. Tissue slices were cut by vibratome (Leica VT1200 S) and recovered for 20 - 30 min at 37 °C before storing at room temperature (25°C) in carbogen-bubbled aCSF before transferral to a recording chamber on the stage of an Olympus BX51W1 microscope maintained at 32 °C. Whole-cell recordings were obtained in granule cells of lobule 9.

### Electrophysiology

Whole-cell patch recordings were obtained using a Multiclamp 700B amplifier, Digidata 1440A to digitize at 40 kHz and analysed with pClamp 10.5 softwea.r Glass pipettes of 1.5 mm O.D. (-AM Systems) were pulled using a-P95 puller (Sutter Instruments; –48 MΩ). For voltage clamp recordings, series resistance was compensated to at least 70% and leak was subtracted offline in pClamp software. Kv4 activation and inactivation plots were recorded from a holding potential of −110 mV stepped in 10 mV (1000 ms) increments to 60 mV, followed by a return step to −30 mV (500 ms) as the test potential. Inactivation curves were fitted according to the Boltzmann equation: *I* /(*I*-*Imax*) = 1/(1 + exp((*Vh*-*V*)/*k*)), where *Vh* is the half inactivation potential and *k* is the slope factor. Activation curves were fitted by the Boltzmann equation: *G* /(*G*-*Gmax*) = 1/(1 + exp((*Va*-*V*)/*k*)), where *G* is calculated with *G* = *I*/(*V*-*Vrev*). *Va* is the half activation potential and *k* is the slope factor. Inactivation and activation plots were constructed using Origin 8.0 (OriginLab, Northampton, MA). ***tsA-201 cells:*** Recordings of Kv4.3 current expressed in tsA-201 cells that express FMRP (WT) or in which *Fmr1* was knocked out by CRISPR-Cas9 (KO) (**Figure 1-2**) were carried out at room temperature with an external solution (pH adjusted to 7.3 with NaOH) composed of (mM): 125 NaCl, 3.25 KCl, 1.5 CaCl_2_, 1.5 MgCl_2_, 10 HEPES, 10 D-Glucose, and 2 TEA. Pipettes were filled with a solution composed of (pH 7.3 with KOH, mM): 110 potassium gluconate, 30 KCl, 1 EGTA, 5 HEPES, and 0.5 MgCl_2_, with 5 di–tris-creatine phosphate, 2 Tris-ATP, and 0.5 Na-GTP added from fresh frozen stock each day. Where indicated, internal levels of calcium concentration were adjusted according to MaxChelator calculations. Previous work (Turner et al., 2016) and unpublished tests on the effects of exchanging internal electrolyte with drugs or a buffered calcium concentration assure complete exchange of the effective internal levels of calcium in 5-10 min of breaking into whole cell recording mode.

### Cerebellar granule cells

Current clamp recordings of cerebellar granule cells from cerebellar sections used an internal solution modified from Gall et al (Gall et al., 2003) composed of (mM): 126 K-gluconate, 4 NaCl, 15 Glucose, 5 HEPES, 1 MgSO_4_, 0.15 BAPTA, 0.05 CaCl_2_, pH 7.3 via KOH, with 5 di–tris-creatine phosphate, 2 Tris-ATP, and 0.5 Na-GTP added from fresh frozen stock each day. Bath solution was carbogen-bubbled aCSF maintained at 32°C. A holding current of less than 10 pA was applied to maintain a holding potential of −70 mV. Input resistance was determined by hyperpolarizing cells with injections of 0 to −8 pA currents in five steps at an increment of −2 pA. Cerebellar granule cell excitability was measured by depolarizing cells from −70 mV with 1 sec steps of currents of 0 - 16 pA in 2 pA increments. Spike events were detected and analysed with pClamp 10.5 software (Axon, Molecular Devices).

### Chemicals and Proteins

Chemical compounds were obtained from Sigma-Aldrich unless otherwise indicated. Drugs were either bath-applied or infused directly through the electrode with the 2PK+ Perfusion system (ALA Scientific, Farmingdale, NY) (Zhan et al., 2020). Drugs were applied at the following concentrations: 30 μM (E)-2-benzylidene-3-(cyclohexylamino)-2,3-dihydro- 1H-inden-1-one (BCI), 0.1 μg/ml MEK1 activated extracellular signal-regulated kinase 1 and 2 (pERK1/2) (SignalChem Biotech), 1 μM TTA-P2 (Alomone Labs), 1 μM Mibefradil (Sigma-Aldrich), and 3 nM Okadaic acid (Sigma-Aldrich). FMRP(1-297)-*tat* was prepared as previously described (Zhan et al., 2020), and was added to the external medium of KO cells for 7 - 8 hr, or delivered *in vivo* by tail vein injection 24 hr before tests (Zhan et al., 2020).

### Immunoprecipitation (IP) and Co-Immunoprecipitation (co-IP)

Protein-protein associations and phosphorylation of the Cav3-Kv4 complex were tested at different levels of [K]o with co-IP and IP as described previously (Zhan et al., 2020). Transfected tsA-201 cells were washed with a low [K]o (LK) solution composed of (mM): 148 NaCl, 1 KCl, 10 HEPES, 3 CaCl_2_, 1 MgCl_2_, 10 Glucose, and equilibrated in LK solution for 10 min. Cells were then exposed to a high [K]o (HK) solution composed of (mM): 148 NaCl, 50 KCl, 10 HEPES, 3 CaCl_2_ 1 MgCl_2_, 10 Glucose for 5 min before being lysed in Lysis Buffer containing (mM): 150 NaCl, 50 Tris, 2.5 EGTA, 1% NP-40, pH 7.5, phosphatase inhibitor (Sigma-Aldrich, Cat. P5726) and proteinase inhibitor (Roche, Cat. 04693124001). Lysates containing 500 µg total protein were incubated at 4°C for 2 - 3 hr with one of the following antibodies: Rabbit anti-Cav3.1 (1:50, custom made and lab verified (Molineux et al., 2006)), rabbit anti- Kv4.3 (1:50, Abcam, Cat. ab65794, RRID:AB_1140929), mouse anti-KChIP3 (1:50, Neuromab, Cat. K66/38, RRID:AB_2877337), rabbit normal IgG (as a control, Abcam, Cat. ab172730, RRID:AB_2687931), mouse normal IgG (as a control, Abcam, Cat. ab37355, RRID:AB_2665484), rabbit anti-P-MAPK substrate (1:50, Cell Signaling Technology, Cat. 14378s, RRID:AB_2798468), rabbit anti-P-MAPK/CDK substrate (1:40, Cell Signaling Technology, Cat. 2325s, RRID:AB_331820). Background control (BC) was used as a negative control by incubating lysate with normal IgG from the same species as the IP antibody. The immuno-complexes were then incubated with 30 μl protein G sepharose beads (GE healthcare, 17-0618-01), which were pre-equilibrated with Lysis Buffer, at 4°C overnight. After four washes with Lysis Buffer, proteins were eluted from sepharose beads by boiling at 95 - 100 °C for 5 min with 40 µl sample buffer diluted from 4x sample buffer (Bio-Rad, 1610747) with Lysis Buffer. IP samples were then resolved and analyzed with Western blot. All commercially obtained antibodies were verified by suppliers. Full western blot data sets are available for viewing on the repository Figshare: https://figshare.com/s/402f8335aa1816cc35c4.

### Western Blot

Cell lysates and IP samples were loaded to 6 - 12% SDS-polyacrylamide gel and resolved with SDS-PAGE. Proteins were transferred to a 0.2 μm PVDF membrane (Millipore) and probed with primary antibodies overnight at 4°C, followed by a goat anti-mouse (1:3000, Invitrogen, Cat.62-6520, RRID:AB_2533947) or donkey anti-rabbit IgG (1:5000, Cytiva, Cat.NA934-1ml, RRID:RRID:AB_772206) HRP-conjugated secondary antibodies. Blot images were taken with a ChemiDoc imager and band densities were analysed with Image Lab (Bio-Rad). Primary antibodies used in this study include: Rabbit anti-FMRP (1:1000, Cell Signalling Technology, Cat.7104s, RRID:AB_10950502), mouse anti-FMRP (1:1000, Abcam, Cat.ab230915, RRID:AB_10950502), rabbit anti-pERK1/2 (1:1000, Cell Signalling Technology, Cat.4370s, RRID:AB_2315112), rabbit anti-tERK1/2 (1:2000, Cell Signalling Technology, Cat.9102s, RRID:AB_330744), mouse anti-GAPDH (1:2000, Invitrogen, Cat.39-8600, RRID:AB_2533438), rabbit anti-DUSP6 (1:1000, Abcam, Cat.ab76310, RRID:AB_1523517), rabbit anti-ɑTubulin (1:2000, Abcam, Cat.ab176560, RRID:AB_2860019).

### Experimental Design and Statistical Analysis

When there were multiple samples in one experiment, samples were randomly chosen to receive different treatments. Experiments were performed with mouse genotypes and experimental groups known to the persons who conducted them. No experimental data without technical issues were excluded. Quantification of Western blot bands was conducted with ImageLab software (Bio-Rad) by normalising a target protein’s intensity to house-keeping proteins. Patch-clamp recordings were analysed with pClamp 10.5 software. All figures were prepared and statistical analyses performed with OriginPro 8 or GraphPad Prism 6, and Adobe Illustrator CC 2021 software. Data normality was tested with the Shapiro-Wilk test. When normally distributed, statistical significance was determined using two-sample Student’s *t*-test for unpaired samples or Welch’s test (*t*-test with Welch correction), and a paired sample Student’s *t*-tests for the same samples under different conditions. When normality was rejected, the Mann-Whitney test was used for two sample comparisons.

## Results

Previous work established that A-type current is subject to modulation by Cav3 T-type calcium current, with a primary effect of modulating the voltage for half inactivation (Vh) of Kv4 channels and thus the availability of A-type current (I_A_) (Anderson et al., 2010a; Heath et al., 2014). The calcium-dependence imparted on Kv4 channels is readily apparent when a reduction of Cav3 channel conductance produces a negative shift in Kv4 Vh, a process that reduces the amplitude of I_A_ (Anderson et al., 2010a, 2010b, 2013; Heath et al., 2014). In the case of cerebellar granule cells the Cav3-Kv4-KChIP3 complex has been shown to increase spike output in relation to long-term potentiation (LTP) of mossy fibre inputs (Heath et al., 2014; Rizwan et al., 2016; Zhan et al., 2020).

FMRP was recently found to exhibit coimmunoprecipitation (co-IP) and FRET with both Cav3.1 and Kv4.3 (Zhan et al., 2020), suggesting a direct association between FMRP and at least these two members of the Cav3-Kv4-KChIP3 complex. Importantly, these protein-protein interactions can be reproduced in tsA-201 cells by co-expressing cDNA for the principal subunits of Cav3.1-Kv4.3-KChIP3 (Anderson et al., 2010a, 2010b, 2013). To further test the molecular interactions between FMRP and subunits of the complex we first transiently transfected cDNA for human Cav3.1, Kv4.3 and KChIP3 in native tsA-201 cells that express FMRP (WT) or cells in which FMRP was knocked out by CRISPR-Cas9 technology (KO) (**Figures 1-2, 1-3**). Kv4.3 Vh was measured using a series of step commands (1 sec) from a holding potential of −90 mV to +60 mV in 10 mV increments followed by a step back to −30 mV (500 msec).

### Cav3.1 and FMRP regulate Kv4.3 availability

We verified the ability for calcium to modulate Kv4.3 Vh by first constructing voltage-inactivation plots for Kv4.3 current in a medium containing 0 calcium (**Fig. 1A**). Addition of 1.5 mM calcium to the bathing medium resulted in a positive shift in Kv4.3 Vh (0 Ca^2+^, −55.5 ± 1.9 mV; 1.5 Ca^2+^, −49.9 ± 1.8 mV; n = 6; *p* = 0.001, paired *t*-test) and an average 142 ± 33.3% (n = 5; *p* = 0.013, paired *t*-test) increase in current amplitude for a step of −40 mV to −30 mV (**Fig. 1A**). Both the calcium-induced shift in Kv4.3 Vh and increase in I_A_ amplitude were blocked by omitting Cav3.1 cDNA from the transfection (**Fig. 1A**). Conversely, applying 1 µM TTA-P2 as a selective Cav3 channel blocker in normal bathing medium containing 1.5 mM calcium negatively shifted Kv4.3 Vh (control, −49.7 ± 1.0 mV; TTA-P2, −54.4 ± 1.7 mV; n = 9; *p* = 0.003, paired *t*-test) without affecting Kv4.3 voltage for activation (Va) (n = 7, *p* = 0.124, paired *t*-test) (**Fig. 1B**). The effects of TTA-P2 on Kv4.3 Vh was further reflected in a 42 ± 7.4% (n = 9; *p* = 4.5 x 10^-4^; paired t-test) decrease in I_A_ amplitude for a step from −40 mV to −30 mV (**Fig. 1B**). The negative shift in Kv4.3 Vh induced by TTA-P2 was comparable to that obtained in 1.5 mM calcium medium for application of either 1 µM mibefradil or 300 µM Ni^2+^, with no significant effects on Kv4.3 Va (**Fig. 1C**) (Anderson et al., 2010a).

**Figure 1.**
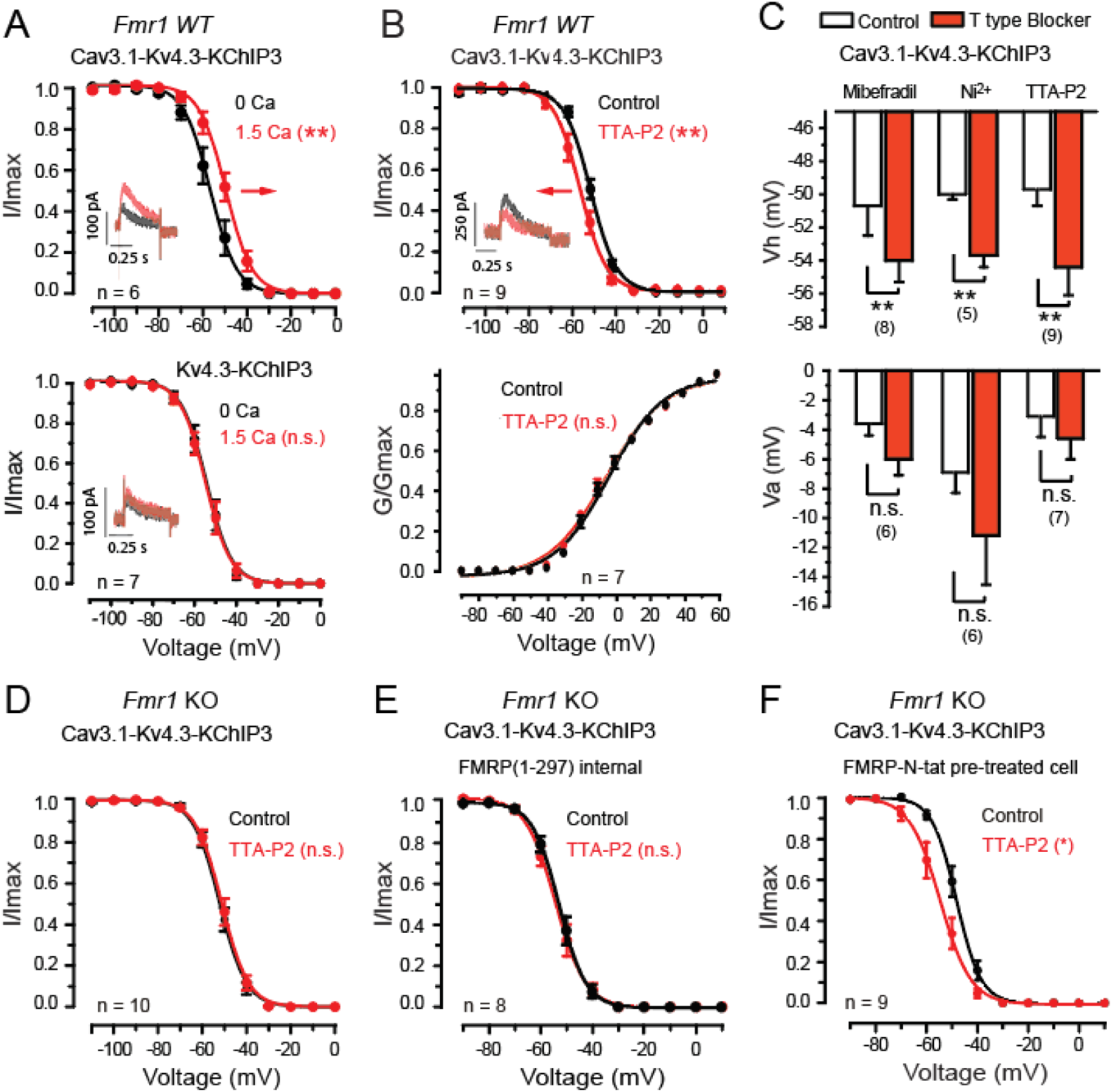
Cav3.1-induced shifts in Kv4.3 Vh are bidirectional and depend on the expression of FMRP. Shown in this and subsequent figures are mean voltage-inactivation or voltage-activation plots for Kv4.3 in *Fmr1* WT cells that express FMRP or in *Fmr1* KO cells in which FMRP was knocked out by CRISPR-Cas9. The proteins coexpressed in each test are indicated above the plots. *Insets* in (**A, B**) show superimposed records of Kv4.3 current evoked by a step from −40 mV to −30 mV before and after the indicated tests. **A,** Raising external calcium from 0 to 1.5 mM induces a positive shift of Kv4.3 Vh (*arrow*) and an increase in Kv4.3 current amplitude that does not occur in the absence of Cav3.1 expression. **B**, Blocking Cav3.1 conductance with 1 µM TTA-P2 induces a negative shift of Kv4.3 Vh (*arrow*), but does not affect the steady state activation of Kv4.3. **C,** Mean bar plots comparing the effects of three Cav3 channel blockers (1 µM mibefradil, 300 µM Ni^2+^, 1 µM TTA-P2) establish the ability to selectively induce a negative shift of Kv4.3 Vh but not Va when applied in a bathing medium containing 1.5 mM calcium. **D,** In *Fmr1* KO cells TTA-P2 (1 µM) fails to shift Kv4.3 Vh. **E, F,** A TTA-P2-induced negative shift in Kv4.3 Vh is not enabled in *Fmr1* KO cells by including 3 nM FMRP(1-297) in the pipette (**E**) but is produced by long term pre-incubation (4 - 8 hr) with 70 nM FMRP-N-*tat* (**F**). Average values are mean ± S.E.M. with sample values (n) indicated on plots. n.s., not significant, ** *p* < 0.01, paired sample *t*-test.

We note that much of the work to date on the Cav3-Kv4-KChIP3 complex was conducted in cerebellar granule cells or WT tsA-201 cells that express FMRP. To test the role of FMRP, we used the *Fmr1* KO cell line. Using KO cells we found that in the absence of FMRP, the ability for TTA-P2 application to induce a negative shift in Kv4.3 Vh in cells was lost (**Fig. 1D**). To further test the effects of a loss of FMRP we recorded Kv4.3 current in KO cells coexpressing the Cav3.1-Kv4.3-KChIP3 complex in standard internal electrode solution or in the presence of 3 nM FMRP(1-297) in the recording electrode. In the presence of internal 3 nM FMRP(1-297), we found no effect of applying 1 µM TTA-P2 on Kv4.3 Vh (**Fig. 1E**). We considered the possibility that reinstating FMRP functions in KO cells might require more time than the 10-15 min typical of whole-cell recordings. It was previously shown that attaching the cell permeating *tat*-peptide moiety to FMRP(1-297) (FMRP-N-*tat*) facilitates its passage across cell membranes (Zhan et al., 2020). We thus pretreated KO cells coexpressing Cav3.1-Kv4.3-KChIP3 with 70 nM FMRP-N-*tat* in the cell medium and found variable levels of recovery at 4 hr, but full recovery at 7 hr. When recordings were subsequently conducted in these cells, the Kv4.3 Vh was not changed between FMRP-N-*tat* pretreated cells to those without this treatment (*p* = 0.07) (**Fig. 1D, F**). However, in cells pretreated for the longer time frame with FMRP-N-*tat*, applying 1 µM TTA-P2 now induced a significant negative shift in Kv4.3 Vh (KO, −48.1 ± 1.4 mV; KO post TTA-P2, −54.8 ± 2.1 mV, n = 9; *p* = 0.012, paired *t*-test) (**Fig. 1F**).

Together these tests are important in providing the first evidence that Cav3 calcium conductance can induce a bidirectional shift in Kv4.3 Vh and I_A_ amplitude in cells expressing the Cav3-Kv4-KChIP3 complex. Moreover, Cav3.1 calcium-dependent modulation of Kv4.3 Vh unexpectedly *depends* on the expression of FMRP, identifying a key role for this molecule in the complex. The ability for FMRP-N-*tat* to rescue calcium-dependent modulation of Kv4.3 in KO cells further indicates that the N-terminal portion of FMRP is sufficient to reinstate the normal functions of the Cav3-Kv4 complex.

### Cav3.1 and KChIP3 are necessary to induce a calcium-dependent modulation of Kv4 Vh

It is known that KChIP molecules can act as the calcium sensor for Kv4 channels with 4 KChIP subunits present in an assembled channel (An et al., 2000; Vierra and Trimmer, 2022) and EF hands 3 and 4 available to bind calcium. Yet the exact role for KChIP3 in Cav3-dependent modulation of Kv4 channels has not been determined.

To test the role of KChIP3 we first coexpressed Cav3.1 and Kv4.3 without KChIP3. In this case applying TTA-P2 (1 µM) produced a very slight shift in Kv4.3 Vh (control, −48.6 ± 1.2 mV; TTA-P2, − 49.7 ± 1.1 mV; n = 10; *p* = 0.003, paired *t*-test) (**Fig. 2A, B**). However, coexpressing KChIP3 with Cav3.1-Kv4.3 restored the ability for TTA-P2 application to induce a much larger negative shift in Kv4.3 Vh (−4.8 ± 3.4 mV, n = 9; *p* = 0.005; two-sample *t*-test) (**Fig. 2B**). The effect of TTA-P2 was lost if an interfering peptide *tat*-pp1 (10 μg/ml) that impedes the Kv4.3-KChIP3 interaction (Scannevin et al., 2004; Callsen et al., 2005) was pre-applied in the bath solution (**Fig. 2B**). We further tested the role of specific EF hands by coexpressing KChIP3 constructs that were mutated to block function in EF hand 3 (KChIP3_(E186Q)_), EF hand 4 (KChIP3_(E234Q)_), or both. Neither of the two single KChIP3 mutants expressed alone with Cav3.1-Kv4.3 fully prevented a TTA-P2-induced negative shift in Kv4.3 Vh (**Fig. 2C**). Coexpressing Cav3.1-Kv4.3 and the double mutant KChIP3_(E186Q, E234Q)_ had no effect on the resting value of Kv4.3 Vh (WT Cav3.1-Kv4.3-KChIP3 Vh, –49.7 ± 1.0 mV, n = 9; WT Cav3.1-Kv4.3-KChIP3_(E186Q, E234Q)_ Vh, −48.5 ± 0.6 mV, n = 8, *p* = 0.334, Welch’s test) but proved to fully block any effects of applying TTA-P2 on Kv4.3 Vh (**Fig. 2C**). These data confirm that calcium sensitivity of the Cav3-Kv4 complex also requires the actions of EF hands 3 and 4 of KChIP3.

**Figure 2.**
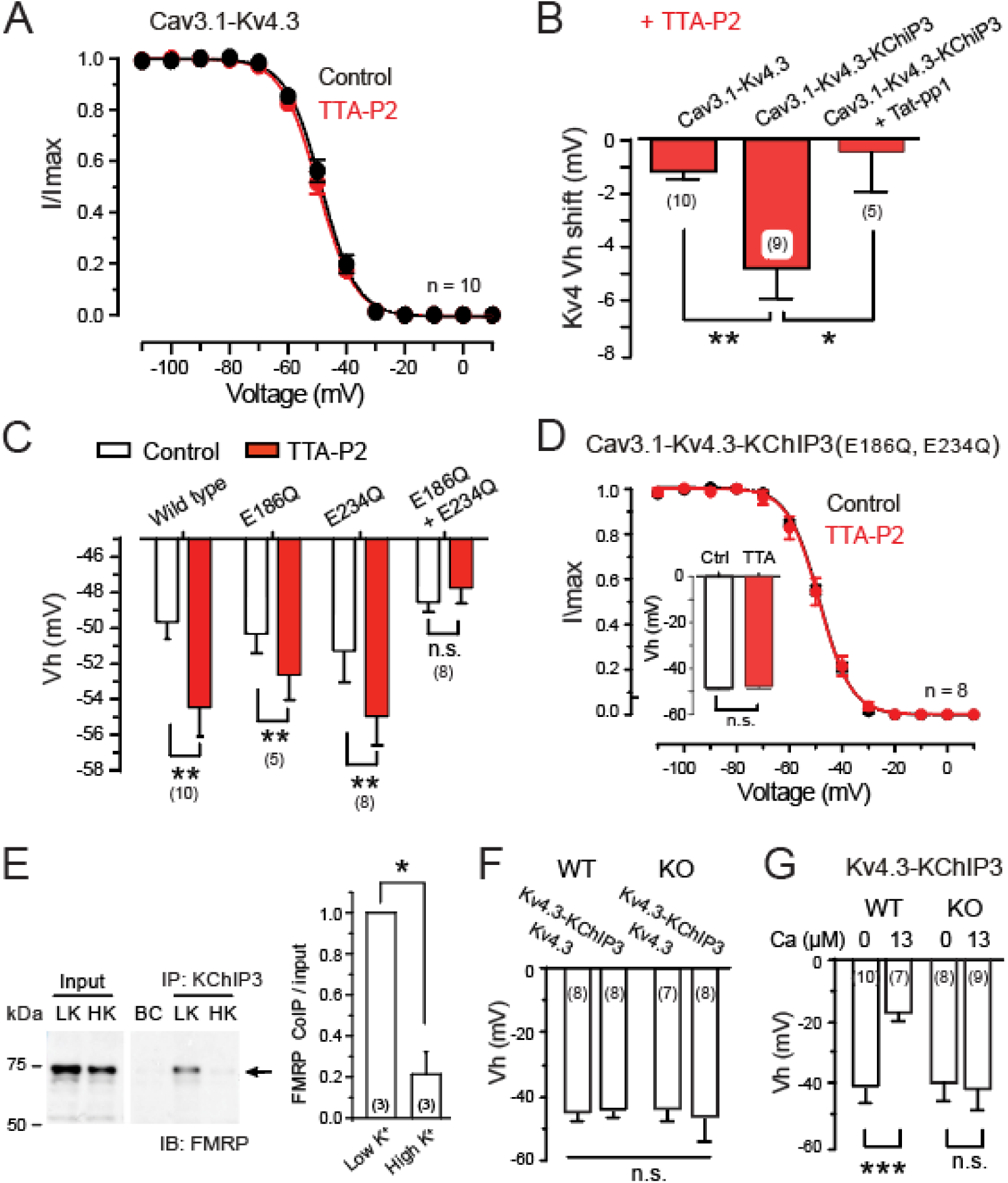
KChIP3 and FMRP are required to enable a calcium- dependent shift in Kv4 Vh. Shown are the effects of TTA-P2 application (**A-D**) on *Fmr1* WT cells coexpressing the indicated subunits. **A, B,** In the absence of KChIP3 coexpression TTA-P2 has only a small effect on Kv4.3 Vh (**A, B**). Coexpressing KChIP3 restores a larger TTA-P2-induced negative shift in Kv4.3 Vh that is lost if preincubated cells with a *tat*-pp1 peptide that interferes with the KChIP3- Kv4.3 binding site (**B**). **C, D,** Mean bar plots showing a TTA-P2-induced shift in Kv4.3 Vh is partially blocked in cells expressing either EF hand mutant, and fully blocked upon expression of a dual KChIP3 EF hand mutant. **E,** Western blot and mean bar plots showing co-IP between KChIP3 and FMRP in *Fmr1* WT cells expressing all subunits of the Cav3-Kv4 complex. The KChIP3-FMRP co-IP detected in the presence of 1.5 mM [K]o (LK) is lost in the presence of 50 mM [K]o (HK) medium to promote Cav3.1 calcium influx. **F,** Coexpressing Kv4.3-KChIP3 in the absence of Cav3.1 has no effect on Kv4.3 Vh in *Fmr1* WT or *Fmr1* KO cells (0 µM internal free calcium, Maxchelator). **G,** Coexpressing Kv4.3-KChIP3 in the absence of Cav3.1 reveals that recording in the presence of 13 µM calcium in the electrode induces a positive shift in Kv4 Vh in *Fmr1* WT cells but not in *Fmr1* KO cells (**G**). BC, background control. Average values are mean ± S.E.M. with sample sizes (n) indicated in brackets. n.s., not significant, * *p* < 0.05, ** *p* < 0.01, *** *p* < 0.001; Paired-sample *t*-test (**C, D**) and Two-sample *t*-test (**B, F, G**).

Previous work established that FMRP forms a link to both Cav3.1 and Kv4.3 (Zhan et al., 2020). To determine if FMRP forms a similar close relationship to KChIP3 as the calcium-sensing component of the complex, we conducted co-IP tests in WT cells. These experiments established that FMRP also co-IPs with KChIP3 (**Fig. 2E; Figure 2-1**). To determine if membrane depolarization and Cav3.1 calcium influx induced any calcium-dependent change in the relationship between KChIP3 and other members of the complex we compared co-IPs obtained in the presence of 1 mM [K]o (Low K; LK) or 50 mM [K]o (High K; HK) solution to promote Cav3.1 calcium influx. HK solution promoted a selective and striking loss of the co-IP between KChIP3 and FMRP (**Fig. 2E**) but not between KChIP3 and either Kv4.3 or Cav3.1 (**Figure 2-1**).

To explore the functional outcome of KChIP3 and FMRP on Kv4 Vh we tested the effects of different levels of internal calcium concentration and with or without FMRP. By coexpressing Kv4.3-KChIP3 in the absence of Cav3.1 and buffering internal calcium concentration to 0 µM (Maxchelator) we determined that adding KChIP3 had no effect on the baseline value of Kv4 Vh in either WT or KO cells (**Fig. 2F**). We then repeated these tests by coexpressing Kv4.3-KChIP3 and recorded with internal solutions that had calcium concentrations buffered to either 0 or 13 µM to compare Kv4 Vh. In WT cells a higher internal calcium concentration induced a significant positive shift in Kv4 Vh (WT 0 calcium, −40.4 ± 1.5 mV, n = 10; 13 µM calcium, −16.9 ± 1.0 mV, n = 7; *p* = 5.3E-9, two-sample *t*-test) (**Fig. 2G**). However, repeating these tests in *Fmr1* KO cells revealed no calcium-dependent shift in Kv4 Vh in the presence of elevated internal calcium (KO 0 calcium, −39.2 ± 2.0 mV, n = 8; 13 µM calcium, −41.2 ± 2.3 mV, n = 9; *p* = 0.64; two-sample *t* test) (**Fig. 2G**), indicating that FMRP is required to invoke a calcium-dependent positive shift in Kv4 Vh.

Together these tests are important in indicating that Cav3 calcium-induced shifts in Kv4.3 Vh and modulation of I_A_ amplitude depends on all of Cav3.1, KChIP3 and FMRP, with a selective and dynamic change in the FMRP-KChIP3 association with an elevation of internal [Ca].

### Cav3.1 and FMRP regulate ERK phosphorylation

We previously found that a negative shift in Kv4 channel Vh in cerebellar granule cells invoked by mossy fibre stimulation could be prevented by PD-98059, an inhibitor of mitogen-activated protein kinase kinase (MEK) that phosphorylates and activates ERK (Rizwan et al., 2016). Yet the factors that control ERK activation and the potential target(s) for ERK phosphorylation of subunits in the Cav3-Kv4-KChIP3 complex are unknown. ERK has been shown to phosphorylate 3 different sites on Kv4.2 that can cause shifts in Vh or Va to regulate I_A_ availability (Adams et al., 2000; Schrader et al., 2002, 2006), but the potential actions of ERK on Kv4.3 have not been reported. Interestingly, Cav3.1-mediated calcium influx has been reported to decrease ERK activation in HEK293 cells (Choi et al., 2005).

To define any Cav3-mediated regulation of the complex by ERK we coexpressed Cav3.1-Kv4.3-KChIP3 in WT cells and used Western blots to detect the effects of membrane depolarization and calcium conductance on ERK1/2 (ERK) activation. For this we used an antibody that targets phosphorylated ERK (pERK) to identify the level of pERK in relation to total ERK (tERK) (**Fig. 3A**). These tests established a baseline ratio of pERK / tERK in low [K]o (1.5 mM), and that application of high [K]o decreased the pERK / tERK ratio by 53.7% within 10 min (high [K]o 0.46 ± 0.08, n = 7, *p* = 4.75E-4, paired *t*-test) (**Fig. 3A**). Moreover, this high [K]o-induced reduction in pERK was blocked by pre-incubation in 1 µM TTA-P2 (high [K]o 0.97 ± 0.28, n = 4, *p* = 0.927, paired *t*-test)(**Fig. 3B**), indicating that a depolarization-induced reduction of pERK depends on Cav3.1 calcium conductance. Repeating these tests in KO cells expressing Cav3.1-Kv4.3-KChIP3 further revealed that the high [K]o-induced reduction in pERK levels was lost in the absence of FMRP (KO high [K]o 0.91 ± 0.10, n = 4, *p* = 0.422, paired *t*-test) (**Fig. 3C**).

**Figure 3.**
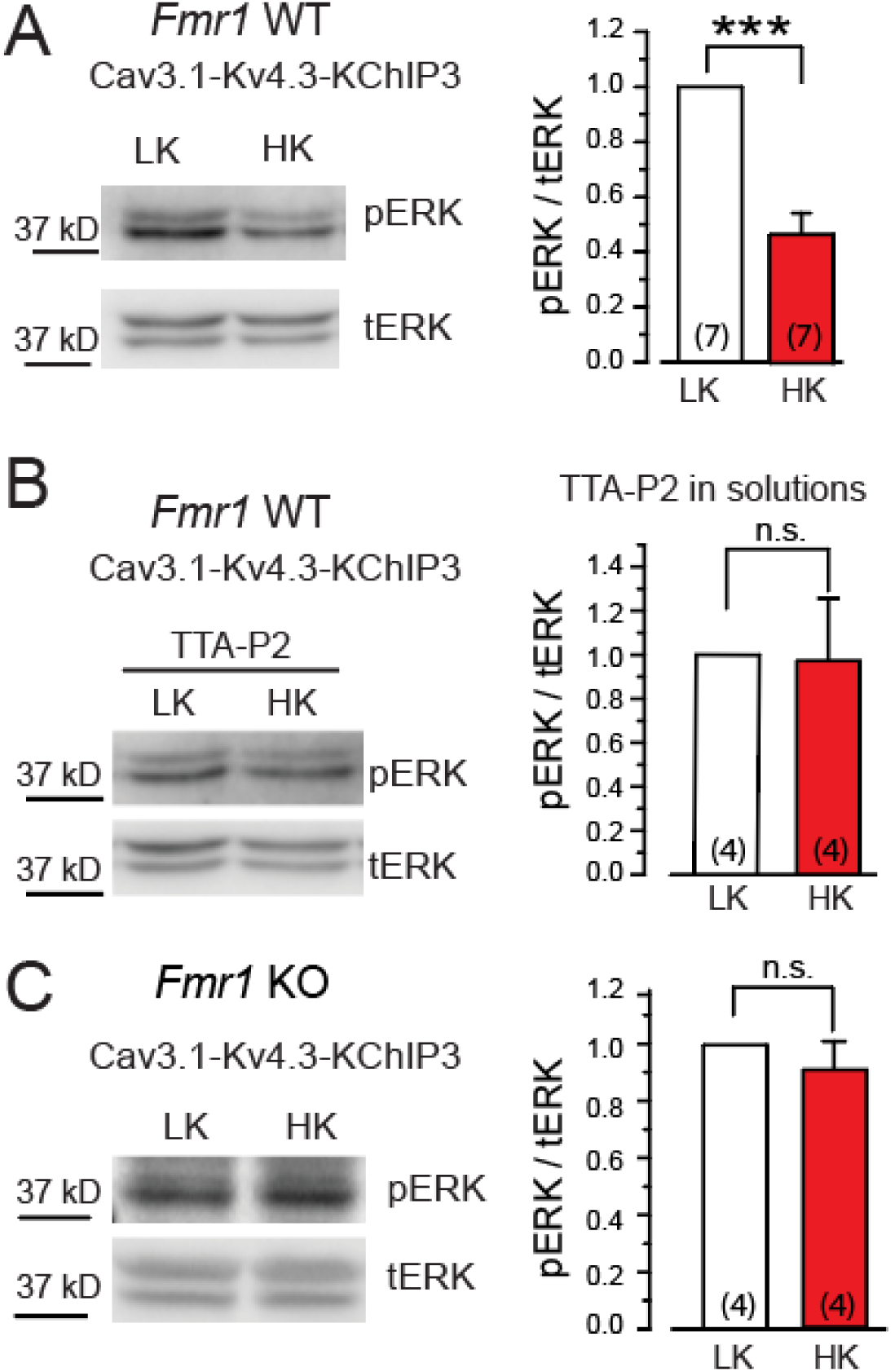
A Cav3.1-dependent reduction in phosphorylated ERK is impaired in *Fmr1* KO cells. Shown are the levels of phosphorylated ERK1/2 (pERK) in relation to total ERK1/2 (tERK) at rest and upon depolarization in tsA-201 cells expressing Cav3.1-Kv4.3-KChIP3. pERK levels are quantified by Western blot and mean bar plots in the presence of 1.5 mM [K]o (LK) or 50 mM [K]o (HK) medium. **A,** In WT cells a resting level of pERK is detected in LK medium. The pERK / tERK ratio is reduced over 10 min upon depolarization in HK medium. **B,** The decrease in pERK / tERK with HK-induced depolarization is prevented if Cav3.1 conductance is blocked by pre-incubation in 1 µM TTA-P2. **C,** The HK-induced reduction of pERK levels is absent in *Fmr1* KO cells. All cells in (**A, B, C**) were also transfected with 0.7µg Kir2.1 to maintain a negative resting potential. Average values are mean ± S.E.M. with sample sizes (n) indicated in bar plots. n.s., not significant, *** *p* < 0.001; Student’s paired sample *t*-test.

These data establish that Cav3.1 calcium conductance lowers the level of activated ERK, and that this process requires the expression of FMRP.

### Cav3.1 activates DUSP to regulate ERK and set Kv4.3 availability

A Cav3.1-mediated reduction in pERK levels could contribute to a shift in Kv4.3 Vh and I_A_ availability. To test this we recorded from WT cells expressing Cav3.1-Kv4.3-KChIP3 with or without 1 μM TTA-P2 in the bath and measured Kv4.3 Vh. For this Kv4.3 current was first measured using a control internal electrolyte that was subsequently replaced by direct electrode infusion with an electrolyte containing 0.2 µg/ml pERK (**Fig. 4A**). When Cav3.1 conductance was blocked in the presence of 1 μM TTA-P2 in the bath, infusion of pERK induced a significant negative shift in Kv4.3 Vh (n = 6, *p* = 0.011, paired *t*-test) (**Fig. 4A**). However, when Cav3.1 conductance was intact (no TTA-P2 in medium), internal infusion of pERK had no significant effect on Kv4.3 Vh (n = 3, *p* = 0.311, paired *t*-test) (**Fig. 4A**). These results are important in suggesting that Cav3.1 calcium conductance can occlude the actions of pERK even when added directly to the electrolyte. To assess the requirement for FMRP in these results we coexpressed the complex in KO cells in the presence of 1 μM TTA-P2 in the bath to prevent calcium conductance, and found that recording with 0.2 µg/ml pERK in the electrode again significantly negative shifted the Kv4.3 Vh (control −45.7 ± 1.8 mV, n = 6; pERK1/2 −53.6 ± 1.5 mV, n = 8; *p* = 0.0069, two-sample *t*-test) (**Fig. 4B**). Together these results indicate the ability for pERK to induce a negative shift in Kv4.3 Vh through a process that is actively prevented or reduced by Cav3.1 calcium conductance.

**Figure 4.**
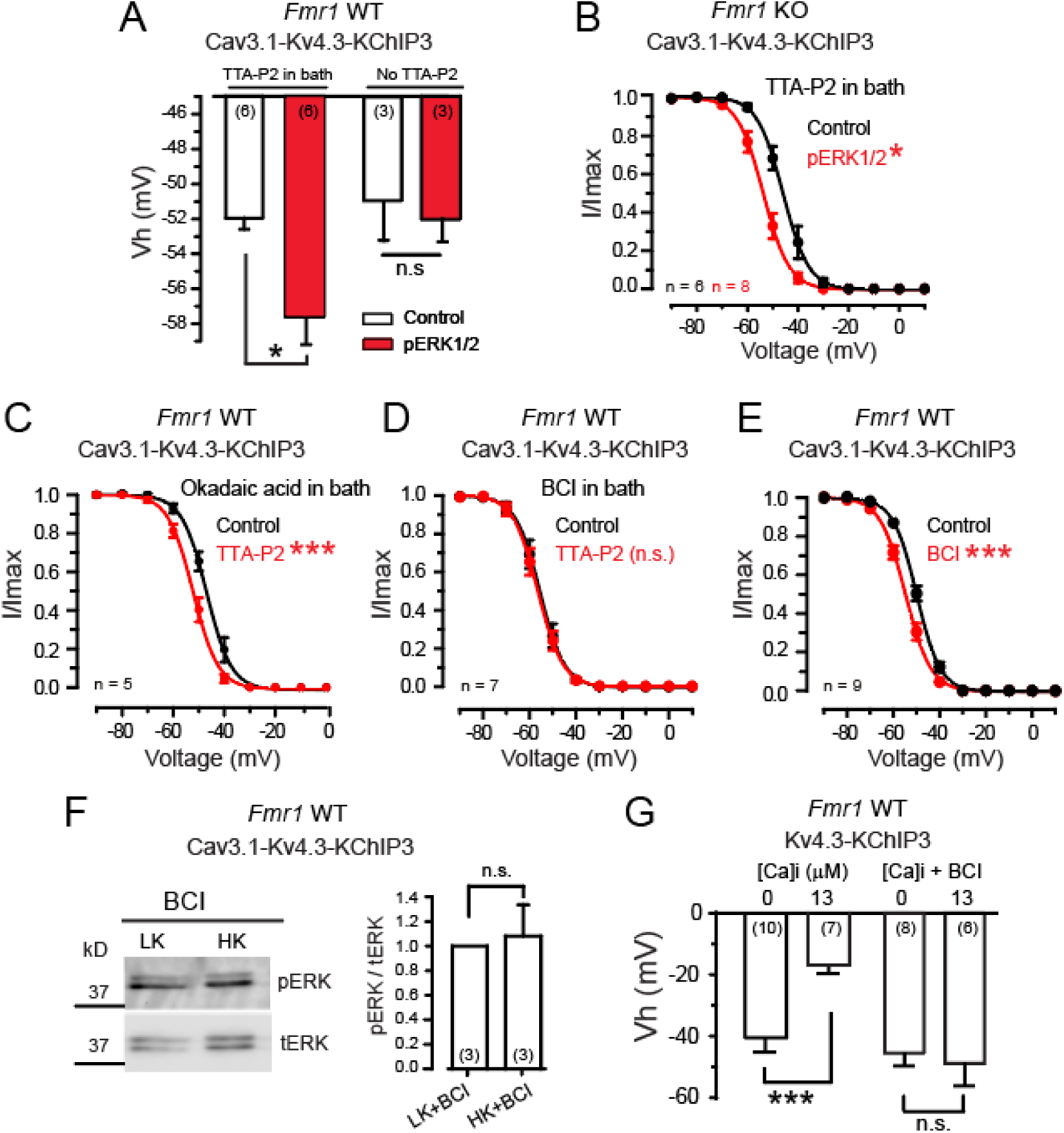
Cav3.1 calcium influx activates DUSP to regulate pERK actions on Kv4.3 Vh. Shown are results from whole-cell recordings and Western blots from *Fmr1* WT or KO cells expressing the indicated subunits. **A, B,** The effects of direct pERK infusion through the electrode in *Fmr1* WT (**A**) or *Fmr1* KO cells (**B**) with Cav3.1- Kv4.3-KChIP3 subunits coexpressed. With 1 µM TTA-P2 in the bath to block T-type current, direct infusion of pERK results in a negative shift in Kv4 Vh in both *Fmr1* WT and KO cells. However, pERK has no effect in the absence of TTA-P2 to retain Cav3.1 conductance (**A**). **C-E,** The effects of phosphatase blockers on Kv4.3 Vh. Bath application of 1 µM TTA-P2 induces a negative shift in Kv4.3 Vh in the presence of 3 nM okadaic acid to block PP1/PP2A phosphatases (**C**), but not in the presence of 30 µM BCI to block DUSP1/6 (**D**). Recordings in the presence of 30 µM BCI also exhibit a negative shift in Kv4.3 Vh (**E**). **F,** Western blots indicating that BCI (30 µM) in the bath blocks a reduction of pERK normally stimulated by a high [K]o-mediated depolarization (HK) in *Fmr1* WT cells expressing the complex (cf. Fig. 3A). **G,** The effects of blocking DUSP1/6 on Kv4.3 Vh in the absence of Cav3.1 expression and direct manipulation of internal calcium concentration in the electrode. Recording with a high level of buffered internal calcium (13 µM, MaxChelator) in the electrode produces a positive shift in Kv4.3 Vh that is blocked in the presence of 30 µM BCI in the bath to block DUSP1/6. Average values are mean ± S.E.M. with sample sizes (n) indicated in bar plots. n.s., not significant, * *p* < 0.05; *** *p* < 0.001; Student’s paired sample *t*- test (**A, C-F**) and Student’s two sample *t*-test (**B, G**).

Cav3.1 calcium influx might reduce pERK levels by activating a phosphatase to dephosphorylate ERK to reduce its actions on downstream targets at serine/threonine (Ser/Thr) sites. The process of dephosphorylation is most often achieved by Ser/Thr phosphatases that include PP1/PP2A (Narayanan et al., 2007). Indeed, Cav3 calcium conductance has been associated with activation of PP2A (Ferron et al., 2011), and PP2A has been shown to dephosphorylate all of FMRP (Narayanan et al., 2007), ERK (Kim et al., 2008) or S6K1 (Hahn et al., 2010). To test the role of PP2A we coexpressed the Cav3.1-Kv4.3- KChIP3 complex in WT cells in the presence of 3 nM okadaic acid to block PP1 and PP2A. We then applied 1 µM TTA-P2 to block Cav3.1 conductance and still found a negative shift in Kv4.3 Vh (control - 46.8 ± 1.4 mV; TTA-P2 -52.2 ± 1.2 mV; n = 5; *p* = 4.98E-5, paired *t*-test) (**Fig. 4C**).

A key factor that specifically controls pERK levels is the family of dual specificity phosphatases (DUSP) that dephosphorylate phospho-threonine and -tyrosine residues on MAPK (Caunt et al., 2008). To test the potential role of DUSPs we coexpressed the Cav3.1-Kv4.3-KChIP3 complex in WT cells and recorded in the presence of 30 µM BCI as an allosteric inhibitor of DUSP1/6 (Molina et al., 2009). Under these conditions perfusion of 1 µM TTA-P2 had no effect on Kv4.3 Vh (control −55.7 ± 1.7 mV; TTA-P2-56.6 ± 1.6 mV; n = 7; *p* = 0.12, paired *t*-test) (**Fig. 4D**), revealing that BCI occluded the effects of Cav3.1 conductance on Kv4.3. In contrast, when Cav3.1-Kv4.3-KChIP3 was coexpressed in WT cells, 30 µM BCI application by itself produced a significant negative shift in Kv4.3 Vh (control −49.8 ± 0.7 mV; BCI - 54.6 ± 1.0 mV; n = 9; *p* = 9.8E-6, paired *t*-test) (**Fig. 4E**). In fact, the BCI-induced negative shift in Kv4.3 Vh was not significantly different from that induced by TTA-P2 (BCI Vh shift, −4.8 ± 0.5 mV, n = 9; TTA-P2 Vh shift, −4.8 ± 1.1 mV, n = 9, *p* = 0.99). These results were important in revealing that a Cav3.1- mediated positive shift in Kv4.3 Vh (cf. **Fig. 1A**) involves Cav3.1 activation of DUSP1/6 and its influence on pERK. This conclusion was further verified using Western blots to repeat the test of depolarizing WT cells using high [K]o medium (cf. **Fig. 3**), which revealed that inhibiting DUSP1/6 with 30 µM BCI prevented a high [K]o-induced depolarization from reducing the pERK/tERK ratio (**Fig. 4F**).

We further explored the extent to which these results could be reproduced by an increase in [Ca]i (**Fig. 4G**). Using whole-cell recordings in WT cells coexpressing only Kv4.3-KChIP3 we buffered [Ca] in the electrode (MaxChelator) to either 0 or 13 µM. Cells recorded in the presence of 13 µM internal calcium exhibited a positive shift in Kv4.3 Vh (**Fig. 4G**). However, no effect on Kv4.3 Vh was detected under these conditions in the presence of 30 µM BCI to block DUSP1/6 (**Fig. 4G**).

These results reveal that a Cav3.1-mediated activation of DUSP1/6 that downregulates the level of pERK is accompanied by a positive shift in Kv4.3 Vh that will increase I_A_.

### Depolarization induces differential effects of phosphorylation of Kv4.3 and FMRP

While the previous data revealed that a Cav3.1-DUSP-ERK signalling pathway regulates calcium dependence of the Cav3-Kv4 complex, the actual targets of ERK-mediated phosphorylation in this process are unknown. The tests already conducted enable predictions on what elements may be important. While regulation of Cav3 channel isoforms by ERK has been reported in fibroblasts and cardiomyocytes (Sharma et al., 2023), we have not detected any change in Cav3.1 density or voltage dependence in relation to stimuli that modulate the Cav3-Kv4 complex (Heath et al., 2014; Rizwan et al., 2016; Zhan et al., 2020). Several lines of evidence implicate Kv4.3 as one important target for phosphorylation. Specifically, shifts in Kv4.3 Vh could be induced in WT cells by direct infusion of pERK in the presence of TTA-P2 to block Cav3.1 conductance (**Fig. 4A**), or with a direct increase in [Ca]i in the absence of Cav3.1 expression (**Fig. 4G**). A negative shift in Kv4.3 Vh could also be induced by internal pERK infusion in WT cells expressing only Cav3.1-Kv4.3 (**Fig. 4A**), suggesting that KChIP3 is not required as an ERK target to enable the calcium-dependent effects studied here. In KO cells coexpressing Cav3.1- Kv4.3-KChIP3 and that lack FMRP, both the calcium-dependent negative shift in Kv4.3 Vh (**Fig. 2G**) and a high [K]o-induced reduction in pERK levels (**Fig. 3C**) were lost, but the effects of pERK on Kv4.3 Vh remained (**Fig. 4C**). These results indicate that pERK actions on Kv4.3 can be sufficient to modulate Kv4.3 Vh but that FMRP is also required for regulating the actions of ERK that promote a calcium-dependent shift in Kv4.3 Vh.

ERK is a member of the MAPK family that can phosphorylate targets at either Serine (PXS*P) or Threonine (PXpTP) motifs (Rubinfeld and Seger, 2005). To identify targets for MAPK-mediated phosphorylation, WT cells coexpressing Cav3.1-Kv4.3-KChIP3 were depolarized using high [K]o medium (HK). Cells were then processed for Western blot analysis for IP between proteins reactive for antibodies specific to MAPK phosphorylated Serine (pSer) or Threonine (pThr) motifs with each of Cav3.1, Kv4.3, KChIP3, and FMRP proteins (**Fig. 5**). These tests revealed pThr sites on FMRP in LK conditions but no detectable difference in phosphorylation levels between LK and HK conditions (**Fig. 5A, B**). No consistent pThr labelling was detected for any of Cav3.1, Kv4.3 or KChIP3 (**Fig. 5A, B**). Tests of immunoprecipitation for Serine motifs revealed pSer labelling for both FMRP and Kv4.3 under resting conditions of LK medium (**Fig. 5C, D**). Exposure of cells to HK medium produced an increase in pSer labelling on FMRP and a marked decrease in phosphorylation of Kv4.3 (**Fig. 5C, D**). Repeating tests for the pSer motif of Kv4.3 further revealed that the reduction of Kv4.3 phosphorylation in HK medium was prevented if cells were preincubated with either 1 µM TTA-P2 (**Fig. 5E**) or 30 µM BCI (**Fig. 5F**). The increase in FMRP phosphorylation was also correlated to Cav3.1 calcium influx although the exact mediator for phosphorylation is unknown since depolarization also lowers the levels of pERK (**Fig. 3A**).

**Figure 5.**
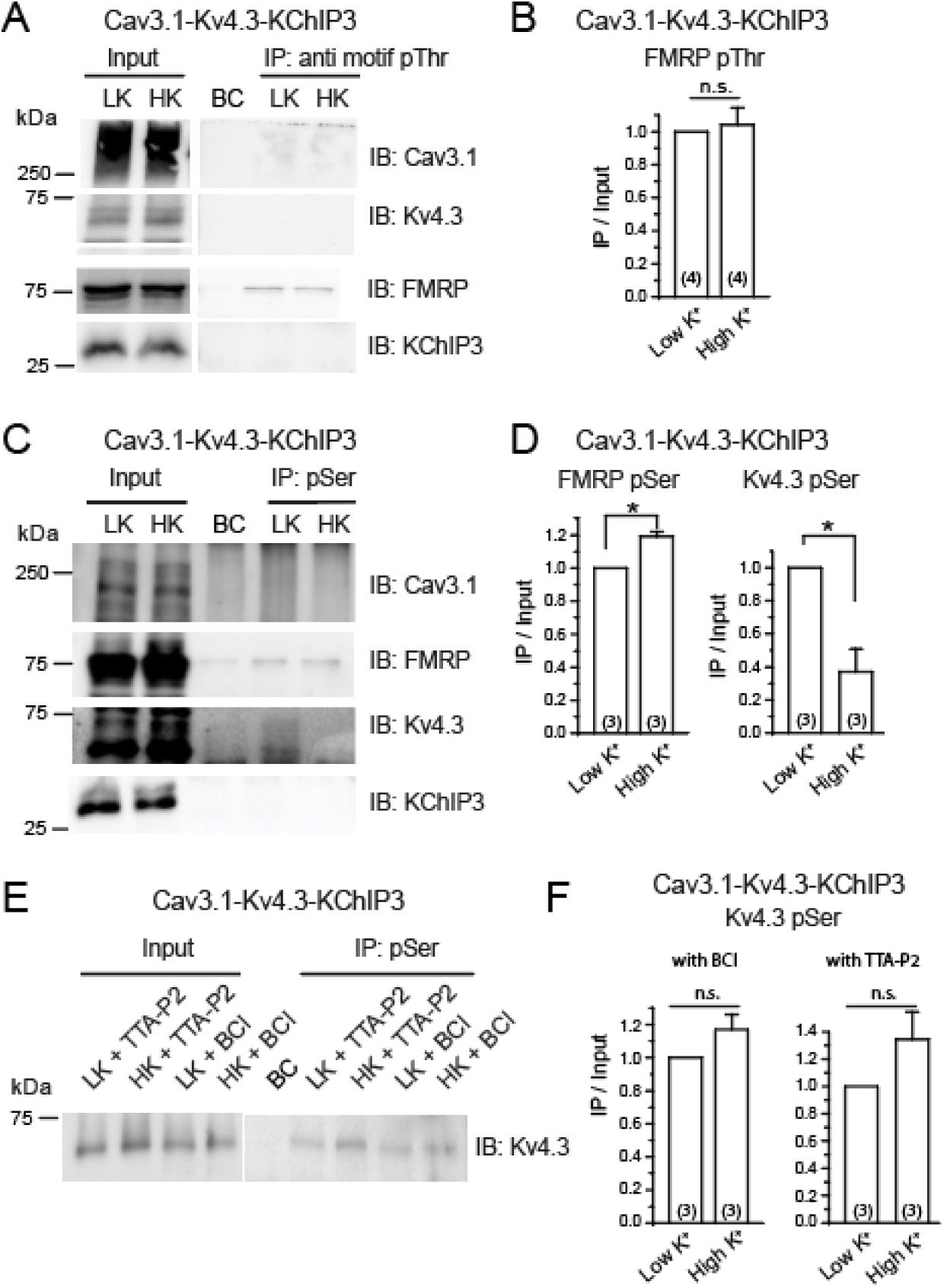
Cav3.1 calcium influx reduces Kv4.3 phosphorylation through a DUSP-ERK pathway. Shown are the effects of testing *Fmr1* WT cells coexpressing Cav3.1-Kv4.3- KChIP3 in either low [K]o (LK) or high [K]o (HK) on phosphorylation of the indicated subunits. Results are shown in Western blots (**A, C, E**) and mean bar plots (**B, D, F**) of the indicated proteins after immunoprecipitation with different ERK substrate antibodies. **A, B,** An anti- PXpTP motif (pThr) antibody detects phosphorylation of FMRP (**A**), but the level of phosphorylated FMRP is not affected by HK treatment (**B**). **C, D,** An anti PXS*P or S*PXR/K motif (pSer) antibody detects phosphorylation of both Kv4.3 and FMRP (**C**), with HK stimulation increasing the level of pSer on FMRP and decreasing the level on Kv4.3 (**D**). **E, F,** Representative western blots (**E**) and mean bar plots of the extent of phosphorylation at pSer sites in LK vs HK conditions (**F**). Pretreating cells with 1 µM TTA-P2 or 30 µM BCI blocks the HK-induced reduction of Kv4.3 phosphorylation. Abbreviations: LK, low potassium; HK, high potassium; BC, background control; IP, immunoprecipitation; IB, immunoblotting. Average values are mean ± S.E.M. with sample sizes (n) indicated in bar plots. n.s., not significant, * *p* < 0.05; Student’s paired sample t-test.

These data reveal that FMRP is phosphorylated at both pSer and pThr sites, and that depolarization increases the pSer labelling. pSer labelling on Kv4.3 is instead reduced upon depolarizing cells, a process that involves Cav3.1 activation of DUSP1/6 and its ability to dephosphorylate ERK. The data thus suggest that Cav3.1 activation of DUSP1/6 and modulation of FMRP and Kv4.3 phosphorylation enable KChIP3 to induce a depolarizing shift in Kv4.3 Vh that increases I_A_ in a calcium-dependent manner (cf. **Fig. 1A**).

### Hyperexcitability of cerebellar granule cells in Fmr1 KO mice is rescued by FMRP-N-tat

Several aspects of excitability of cerebellar granule cells and their response to synaptic input have been shown to depend on the Cav3.1-Kv4-KChIP3 complex (Heath et al., 2014; Rizwan et al., 2016; Zhan et al., 2020). To further test the effects of FMRP-N-tat on excitability of neurons we recorded from lobule 9 granule cells from P22 - 24 WT and *Fmr1* KO mice in cerebellar slices prepared 1 day following tail vein injection of either vehicle or 1 mg/kg FMRP-N-*tat*. Whole-cell recordings were used to deliver 1 sec current pulses to map the current-frequency relationship of sodium spike discharge. In WT mice injected with vehicle alone, granule cells fired with an initial frequency of 0.5 ± 0.54 Hz, showing a linear increase in firing with increments of current injection up to 42.2 ± 8.7 Hz (n = 9) (**Fig. 6A, B**). The maximum firing frequency reached during 16 pA current injection increased by 86% in *Fmr1* KO granule cells compared to WT cells, further indicating a state of hyperexcitability (**Fig. 6C**). An increased sensitivity to depolarizations was also apparent in a decrease in the rheobase to evoke spike firing in vehicle injected *Fmr1* KO mice compared to WT mice (**Fig. 6D**). Finally, the increased spike frequency was associated with a significant decrease in the decay tau (rate of repolarization) of evoked spikes **(Fig. 6E**). By comparison there was no significant difference in the input resistance of granule cells between WT and *Fmr1* KO mice (WT 1.05 ± 0.10 GΩ, n = 9; *Fmr1* KO 1.30 ± 0.17 GΩ, n = 7; *p* = 0.24, two-sample *t*-test). We then injected *Fmr1* KO mice with 1 mg/kg FMRP-N-*tat* and prepared tissue slices 24 hr later, a time previously shown to be sufficient for this compound to distribute throughout the cerebellum for uptake by granule and Purkinje cells (Zhan et al., 2020). Here we found complete rescue of the elevated firing rate of *Fmr1* KO compared to WT granule cells over the full range of a current-frequency plot, with similar effects on rheobase and rate of spike repolarization (**Fig. 6A-E**). These effects by FMRP-N-tat are important in that IA has previously been shown in granule cells to shape each of the parameters tested here (Heath et al., 2014; Zhan et al., 2020).

**Figure 6.**
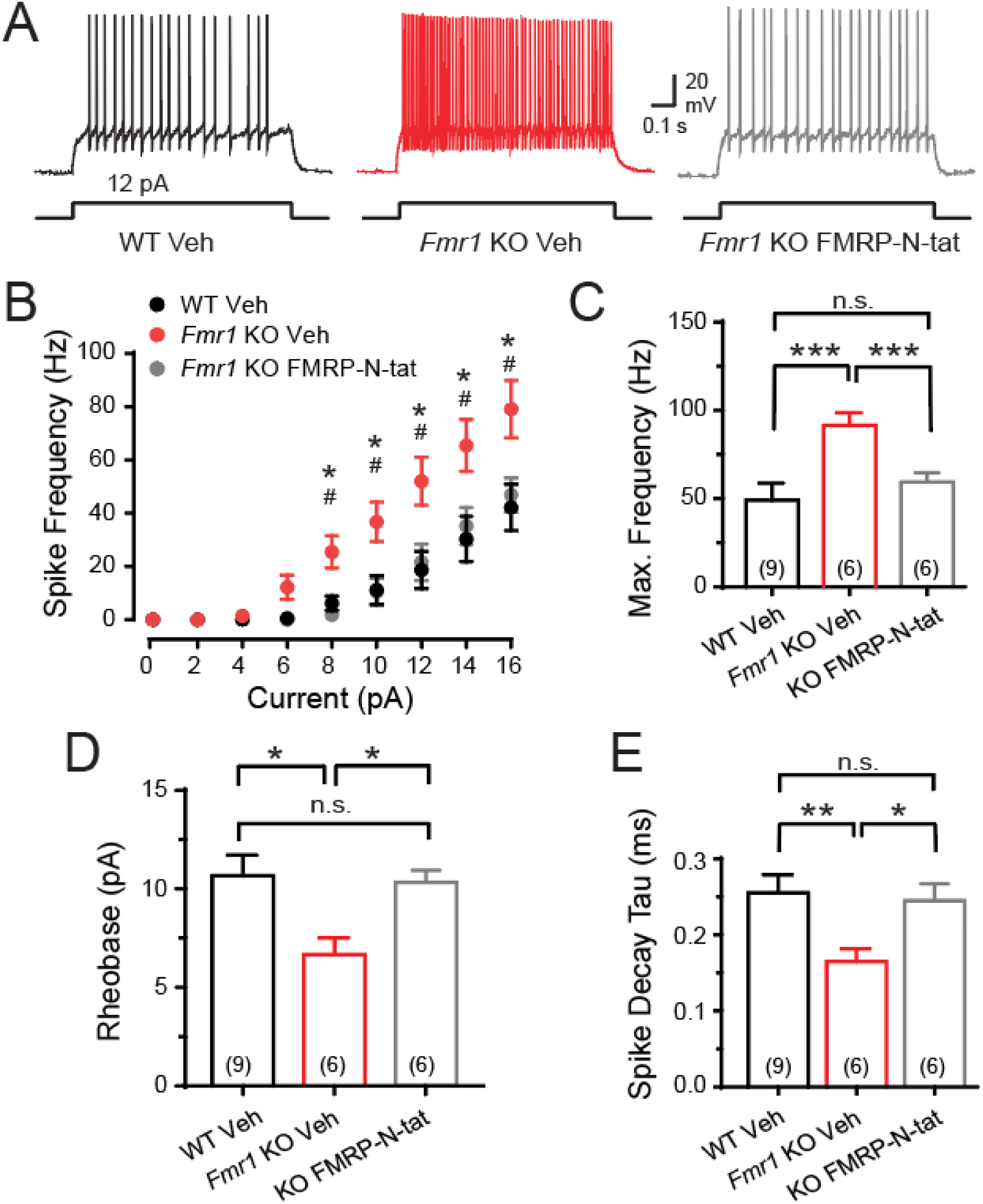
FMRP regulates excitability in cerebellar granule cells. The effects on spike output of cerebellar granule cells in tissue slices of P22 -24 male WT and *Fmr1* KO mice 24 hr following tail vein injection of vehicle or 1.0 mg/kg FMRP-N-tat. **A**, Representative whole-cell recordings indicate that tonic firing frequency in response to current injection is markedly elevated in vehicle injected *Fmr1* KO mice compared to WT mice. Firing frequency is rescued in granule cells of *Fmr1* KO mice 24 hr after FMRP- N-tat injection. **B,** Mean frequency-current (F/I) plots of spike output for 1 sec long current injections. **C**, Bar plots of mean maximal firing frequency measured from the first 4 spike outputs with 16 pA current injection. **D**, Bar plots of rheobase current required to reach spike firing threshold. **E**, Bar plots of mean decay tau of spike output as a measure of the rate of repolarization with the rheobase current injection required to trigger spike firing. Average values are mean ± S.E.M. with sample sizes (n) indicated in brackets. n.s.; not significant; * *p* < 0.05, ** *p* < 0.01, and *** *p* < 0.001 (**C, D, E**); # *p* < 0.05 for *Fmr1* KO vehicle vs FMRP-N-tat injected firing rate (**B**); Student’s two sample *t*-test.

### An imbalance in expression levels of DUSP in Fmr1 KO cells is rescued by FMRP-N-tat

The function of the Cav3.1-DUSP-ERK signalling pathway here was defined primarily using WT or KO tsA-201 cells. To explore the prevalence of DUSP6 expression *in situ*, we conducted immunocytochemical labelling for DUSP6 in the cerebellum of P60 male mice. These tests confirmed a widespread expression of this phosphatase in granule cells (**Fig. 7A**). The DUSP family is interesting in establishing a reciprocal inhibitory system with pERK that will activate nuclear transcription factors that modify DUSP levels (Rubinfeld and Seger, 2005; Cagnol and Rivard, 2013; Caunt and Keyse, 2013). We used Western blot analysis to explore the potential for a loss of FMRP to affect the levels of DUSP protein. This was first tested in WT and KO tsA-201 cells exposed in culture medium to either vehicle alone or FMRP-N-tat (70 nM) for 7 hr (**Fig. 7B**). These tests revealed a significant decrease in the level of DUSP6 in KO cells compared to WT cells. However, the lower levels of DUSP6 in KO cells were rescued by pretreating cells with 70 nM FMRP-N-tat for 7 hr (**Fig. 7B**), a timeline that proved to overlap that required for FMRP-N-tat to rescue Cav3.1 modulation of Kv4.3 in *Fmr1* KO cells (cf. **Fig. 1F**).

**Figure 7.**
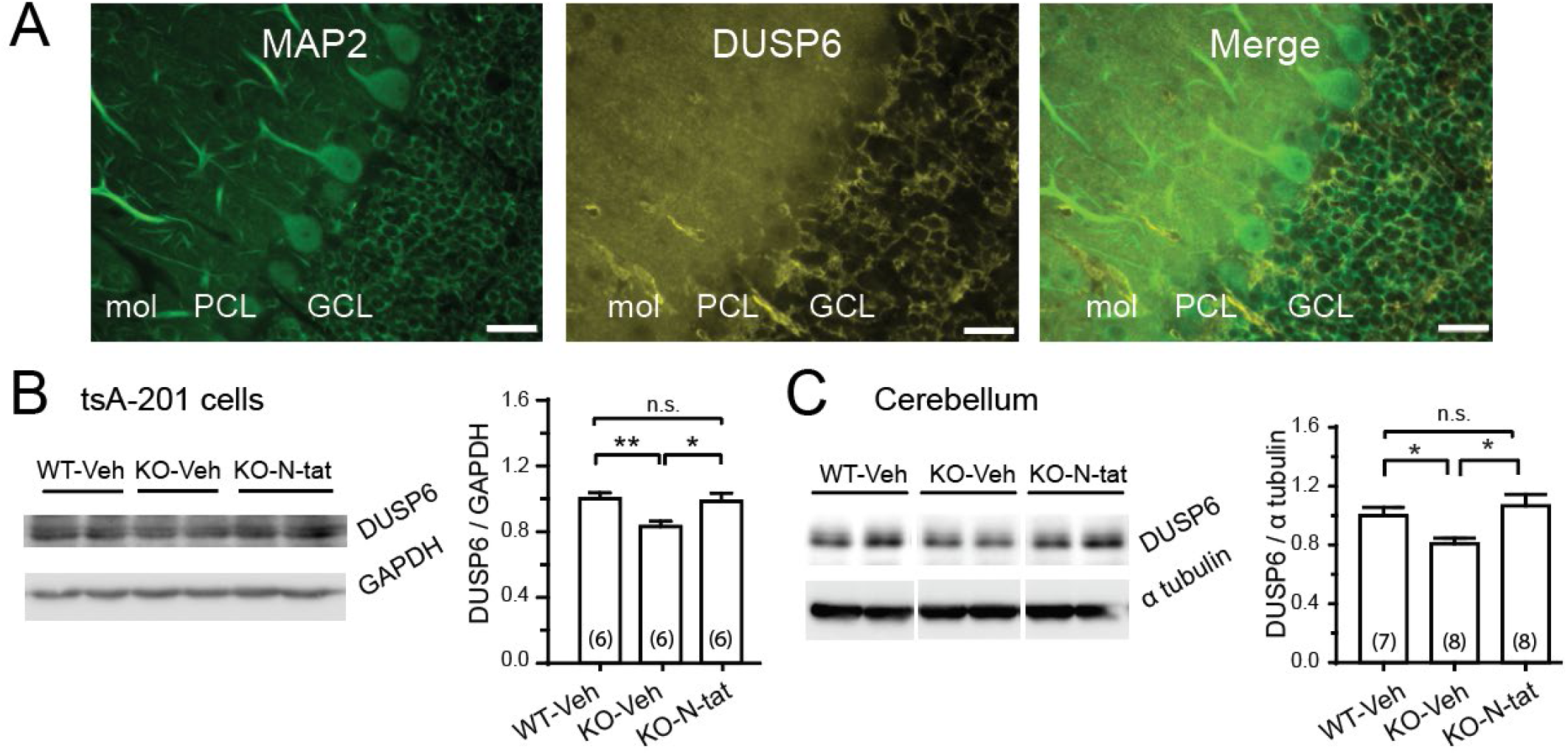
DUSP6 expression in cerebellum is disrupted in *Fmr1* KO cells but rescued by reintroducing FMRP-N-tat. **A,** Low power immunolabel images for MAP-2 as a structural label and DUSP6 in tissue sections of P60 WT mouse lobule 9 cerebellum showing DUSP6 expression in granule cells. **B, C,** Two representative Western blots of the relative expression levels of DUSP6 in homogenates of *Fmr1* WT and KO tsA-201 cells (**B**) and P24-26 mouse cerebellar WT and *Fmr1* KO tissue (**C**). Protein levels are normalized to levels of GAPDH (**B**) and 𝛂𝛂-tubulin (**C**). DUSP6 levels are lower in tsA-201 *Fmr1* KO cells (**B**) and *Fmr1* KO cerebellum compared to WT (**C**). Direct application of 70 nM FMRP-N-tat in tsA-201 culture cell medium restores the level of DUSP6 (**B**), as found in *Fmr1* KO cerebellar tissue 24 hr after tail vein injection of 1.0 mg/kg FMRP-N-tat (**C**). Average values are mean ± S.E.M. with sample sizes (n) indicated in bar plots. n.s., not significant, * *p* < 0.05, ** *p* < 0.01. Student’s two sample *t*-test. Mol, molecular layer; PCL, Purkinje cell layer; GCL, granule cell layer. Scale bars (**A**), 50 µm.

We then compared these results to the levels of DUSP detected in homogenates of whole cerebellum from WT and *Fmr1* KO mice 24 hr following tail vein injection of either vehicle or FMRP-N-tat (**Fig. 7C**). In this case we found that the level of DUSP6 in cerebellum was significantly reduced in *Fmr1* KO mice compared to WT mice (**Fig. 7B**). Yet once again, tail vein injection of *Fmr1* KO mice with 1 mg/kg FMRP-N-tat promoted a rescue of the level of DUSP6 in the cerebellum when tested 24 hr later (**Fig. 7C**).

These data are important in identifying dual actions of FMRP-N-tat in reassociating with the Cav3- Kv4 complex at the plasma membrane to restore calcium-dependent regulation (**Fig. 6**) as well as restoring a disrupted level of DUSP (**Fig. 7**) that is critical to the function of the Cav3-Kv4 complex.

## Discussion

Cav3 calcium channels form a nandomain association with a Kv4-KChIP complex to confer calcium-dependent modulation of I_A_ that can influence signal processing (Vierra and Trimmer, 2022). Despite our knowledge of the functional outputs of the Cav3-Kv4 complex, the molecular mechanisms underlying calcium-dependent control of Kv4 channels had not been fully explored. The current study identified a critical role for FMRP in enabling a new Cav3-DUSP signalling pathway to regulate ERK-mediated phosphorylation in the complex. The physiological relevance of FMRP to Cav3-Kv4 function was further shown by the rescue of a hyperexcitable state and DUSP levels in cerebellar granule cells in the *Fmr1* KO mouse model for 24 hr by *in vivo* administration of FMRP-N-tat.

### FMRP as a member of the Cav3-Kv4-KChIP complex

An association between Kv4 potassium channels and KChIP proteins as a potential calcium-sensing element has long been recognized (An et al., 2000; Jerng et al., 2005; Pioletti et al., 2006; Wang et al., 2007), followed by recognition of Cav3 calcium channels as a component of the complex (Molineux et al., 2005). The role of FMRP as a critical element tightly associated with this complex was identified through co-IP or FRET imaging. Thus, FMRP co-IPs with each of Cav3.1, Kv4.3, and KChIP3 (**Figs. 2, 2-1**) and exhibits FRET with both Cav3.1 and Kv4.3 (Zhan et al., 2020). The FMRP-KChIP3 association proves to be unique in exhibiting a calcium-dependent loss of co-IP (i.e. dissociation) (**Fig. 2**), a result not apparent for the co-IP between FMRP and either Cav3.1 or Kv4.3 (**Figure 2-1**). The current work using a *Fmr1* KO tsA-201 cell line provides evidence that modulation of Kv4.3 also involves Cav3.1 calcium-dependent activation of DUSP to control the levels of pERK-mediated phosphorylation. It is interesting to note that several studies have been carried out on how the Cav3-Kv4 complex shapes the postsynaptic response of cerebellar stellate and granule cells (Molineux et al., 2005; Anderson et al., 2010a, 2010b, 2013; Heath et al., 2014; Rizwan et al., 2016; Zhan et al., 2020). A retrospective analysis indicates that the new findings of a role for FMRP in the Cav3-Kv4 complex do not conflict in any way with results of these studies. Rather, FMRP proves to have been a hidden component that is central to the entire process by which Kv4 channels achieve calcium-dependent regulation.

### Calcium-dependent control of Kv4 and I_A_ availability

Our available data supports a working model of actions that underlie calcium-dependent regulation of Kv4.3 for the combination of FMRP, Cav3.1, Kv4.3 and KChIP3 (**Fig. 8**). Under normal conditions FMRP is associated with each of the protein subunits of the Cav3.1-Kv4.3-KChIP3 complex, with some resting level of phosphorylation of both Kv4.3 and FMRP at MAPK consensus sites that maintains a relatively low amplitude I_A_ (**Fig. 8A**). Membrane depolarization induces Cav3.1-mediated calcium entry that activates DUSP1/6 to dephosphorylate ERK. The decrease in pERK levels is then reflected in a reduction in phosphorylation of Kv4.3 along with an increase in phosphorylation of FMRP. Associated with calcium influx is a dissociation of FMRP-KChIP3, and a KChIP3-induced positive shift in Kv4.3 Vh that increases I_A_ amplitude to reduce membrane excitability (**Fig. 8B**). The converse effect of a hyperpolarizing shift in Kv4.3 Vh can be revealed by reducing Cav3.1 calcium conductance, decreasing [Ca]i, or by direct intracellular infusion of pERK. In the *Fmr1* KO mouse cerebellum, the level of DUSP6 is reduced, indicating a role for FMRP in regulating translation and expression of a phosphatase important to Cav3-Kv4 function (**Fig. 8C**). The lack of FMRP further removes the ability to phosphorylate FMRP and its dissociation from KChIP3, which together blocks all calcium-dependent regulation of Kv4.3 Vh and I_A_ amplitude (**Fig. 8C**). Thus it appears that FMRP is required for KChIP3 to carry out its role of producing calcium-dependent shifts in Kv4.3 Vh. This model helps account for the sensitivity of Kv4 channels to fluctuations of calcium conductance that provides a bidirectional control over subunit phosphorylation by ERK and IA availability.

**Figure 8.**
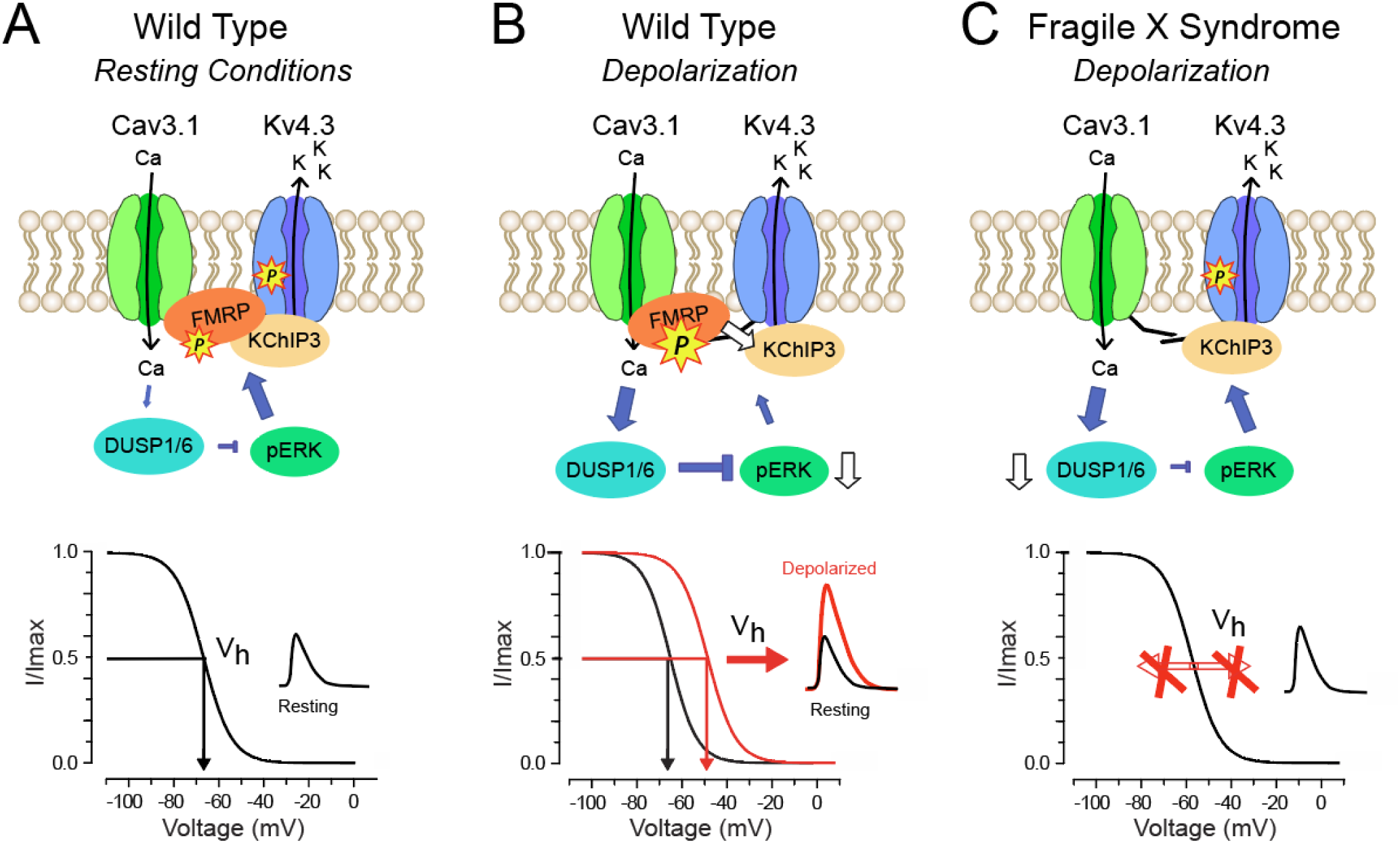
Model of FMRP regulation of calcium-dependent modulation of Kv4 channels via a DUSP-ERK pathway. Interactions shown are relevant to mouse cerebellar lobule 9 granule cells where Cav3.1, Kv4.3, KChIP3 and FMRP form a tight association at the plasma membrane. **A-C,** Shown are interactions found at rest and in response to depolarization in WT cells (**A, B**), and the effects of depolarization in the case of a loss of FMRP in FXS (**C**). **A,** Under resting conditions, a low Cav3.1 conductance provides little activation of DUSP1/6 that leads to an elevation of pERK. Both FMRP and Kv4.3 exhibit phosphorylation at MAPK consensus sequence sites, promoting a hyperpolarizing shift in Kv4.3 Vh (*white arrow*) that reduces IA amplitude. **B,** Upon membrane depolarization Cav3.1 calcium conductance increases, activating DUSP1/6 to reduce levels of pERK and MAPK phosphorylation of Kv4.3 at a pSer site, and an increase in phosphorylation at a pSer site on FMRP. Accompanying this is a reduction of the FMRP-KChIP3 association (*white arrow*) and a depolarizing shift in Kv4.3 Vh (*red filled arrow*) that increases IA amplitude. **C,** In Fragile X Syndrome a loss of FMRP from the complex and its interactions with KChIP3 prevents all calcium-dependent regulation of Kv4.3 Vh and IA amplitude (double arrows marked by X). In the *Fmr1* KO cerebellum the level of DUSP6 is reduced but rescued along with function of the Cav3-Kv4 complex upon reintroducing the FMRP(1-297) fragment as a tat- conjugated peptide.

### Distinct pathways to modulate I_A_ and membrane excitability

It is well established that mGluR/NMDAR activation provides a source of calcium influx to trigger the interplay between ERK, PP2A and S6K1 to modify the phosphorylation state of FMRP (Bagni and Greenough, 2005; D’Incal et al., 2022). Bursts of mossy fiber input to stimulate mGluR/NMDARs on granule cells was also shown to activate ERK that reduced I_A_ to increase spike output as part of mossy fiber LTP (Rizwan et al., 2016). Yet the current work reveals a new Cav3.1 channel-activated signalling pathway that does not involve PP2A and instead deactivates ERK to increase I_A_.

The data thus suggest the existence of two pathways that can separately employ ERK to modulate I_A_ availability. One is intrinsic to the cell in being triggered by Cav3.1 calcium influx to activate DUSP and reduce pERK and phosphorylation of Kv4.3. This Cav3-dependent process also depends on a dynamic change in an FMRP-KChIP3 association, with the net effect of inducing a depolarizing shift of Kv4.3 Vh to increase I_A_ and reduce spike frequency. The second pathway requires an mGluR/NMDAR stimulus that activates ERK to induce a hyperpolarizing shift in Kv4 Vh to decrease I_A_ and promote a long term increase in excitability in a manner that is also FMRP-dependent. At this time the exact means by which mGluR/NMDA activation results in a pERK-mediated increase in excitability is unknown but is expected to involve both pre- and postsynaptic mechanisms (Zhan et al., 2020). However, both the intrinsic and synaptically evoked process depend on calcium-dependent modulation of Kv4 Vh that in turn relies on FMRP.

### Rescue of FMRP function in FXS

A number of strategies have been developed to reduce the symptoms of FXS through pharmacology (Yamasue et al., 2019), reactivating FMRP expression (Xie et al., 2016; Graef et al., 2019; Yrigollen and Davidson, 2019; Shah et al., 2023), or inducing FMRP expression through delivery of viral constructs (Hampson et al., 2019). It was recently found that reintroducing the N-terminus of FMRP as a tat conjugate peptide achieved rapid transport across the BBB, rescuing circuit functions and alleviating symptoms of *Fmr1* KO mice (Zhan et al., 2020). The current work now shows that a loss of FMRP and its actions in the Cav3-Kv4 complex increases the intrinsic excitability of cerebellar granule cells to promote a hyperexcitable state of postsynaptic spike discharge. Tail vein injection of FMRP-N-tat fully rescued granule cell excitability in terms of postsynaptic spike properties and the level of DUSP even 24 hr after tail vein injection. It is striking that FMRP-N-tat is able to restore the actions of FMRP both within an ion channel complex at the membrane while also modifying the expression level of a phosphatase that is key to calcium-dependent modulation of IA. It is important to note that the N-terminal region of FMRP has also been shown to restore GABAergic inhibition from basket cells to Purkinje cells in *Fmr1* KO mice (Yang et al., 2018). Conditional expression of FMRP in Purkinje cells of global *Fmr1* KO mice further restores Purkinje cell excitability (Stoodley et al., 2017; Gibson et al., 2023). The potential thus exists for FMRP fragments to exert therapeutic actions on several different parameters of cerebellar circuit dysfunction in FXS.

### Cerebellar role in ASD in FXS

The behavioural and cognitive functions affected in FXS fall into the realm of ASD. The cerebellum is highly implicated given its established role in higher levels of cognition (D’Mello and Stoodley, 2015; Stoodley et al., 2017; Mapelli et al., 2022; Gibson et al., 2023). Indeed, a change in the output of cerebellar Purkinje cells in the right Crus I/II region has been shown to account for a range of ASD symptoms that include hypersensitivity to sensory stimuli (Tsai et al., 2012, 2018; D’Mello and Stoodley, 2015; Cupolillo et al., 2016; Peter et al., 2016; Stoodley et al., 2017; Sundberg et al., 2018; Gibson et al., 2023). A loss of FMRP has also been shown to alter the activity of granule (Zhan et al., 2020), Purkinje (Koekkoek et al., 2005; Gibson et al., 2023; Martín et al., 2023), and basket cells (Yang et al., 2018). Our work was conducted on granule cells in the vermis of lobule 9 in the posterior cerebellum, a region that has a role in processing multiple sensory inputs and exhibits functional connectivity with limbic structures (Sang et al., 2012; Markwalter et al., 2019). To our knowledge a direct link between granule cell activity and ASD symptoms in FXS has not yet been defined as for Purkinje cells and ASD. Yet, a change in the intrinsic excitability of granule cells will alter signal processing at the first stage of mossy fiber input to the cerebellum that will affect cerebellar cortical activity, and finally the output to neocortical regions that are predicted to contribute to ASD.

## Author Contributions

X.Z. and R.W.T. designed the experiments, X.Z. acquired and analysed electrophysiology and protein biochemical data, P.P. designed and provided KChIP3 EF hand mutant cDNA, and H.A. contributed protein biochemistry. R.W.T. and X.Z. supervised the study and wrote the manuscript. R.W.T. and X.Z. have direct access and verified all data reported in the manuscript. All authors approved the final version of the submitted manuscript.

## Acknowledgments

This work was supported by the Canadian Institutes for Health Research and Fragile X Canada / FRAXA, and a SFARI Explorer grant. Postdoctoral Fellowships (X.Z.) provided by Fragile X Canada / FRAXA (X.Z.), the Hotchkiss Brain Institute, and the Cumming School of Medicine (Calgary). We gratefully acknowledge J. Forden, F. Visser, and Y. Yu for expert technical assistance and Dr. N. Cheng for comments on the manuscript.

## Extended Data

**Figure 1-1.**
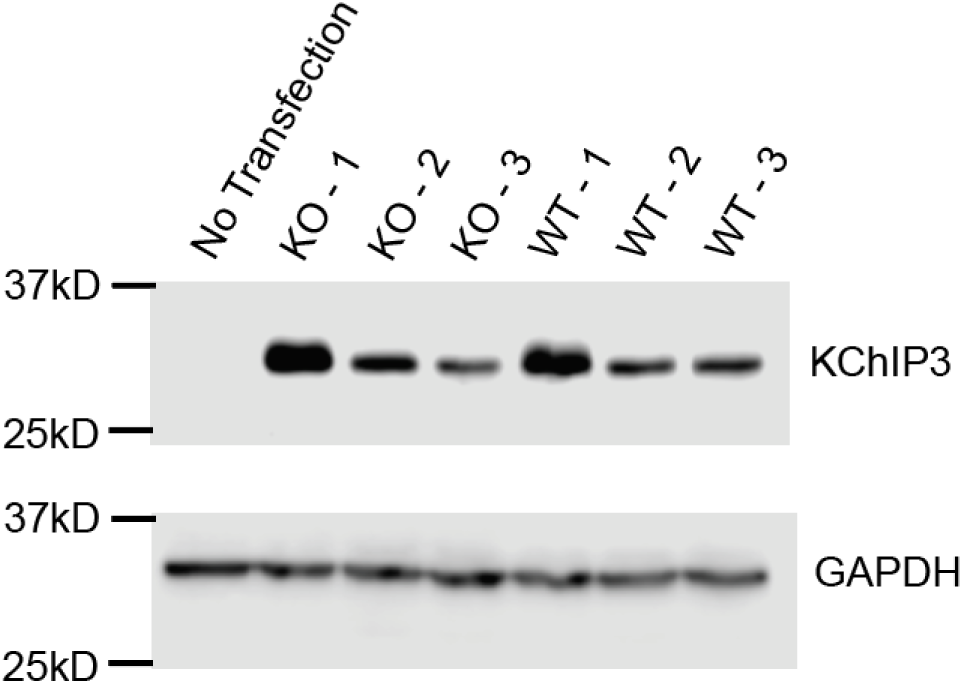
KChIP3 is not endogenously expressed in tsA-201 cells. Shown is a Western blot of WT or KO tsA-201 cell lysates probed with an anti-KChIP3 antibody. First lane, lysate from non-transfected tsA- 201 cells; Lanes 2 - 4, lysates from three different batches of *Fmr1* KO tsA-201 cells transfected with Cav3.1-Kv4.3-KChIP3; Lanes 5 - 7, lysates from three different batches of WT tsA-201 cells transfected with Cav3.1-Kv4.3-KChIP3. GAPDH was used as a loading control.

**Figure 1-2.**
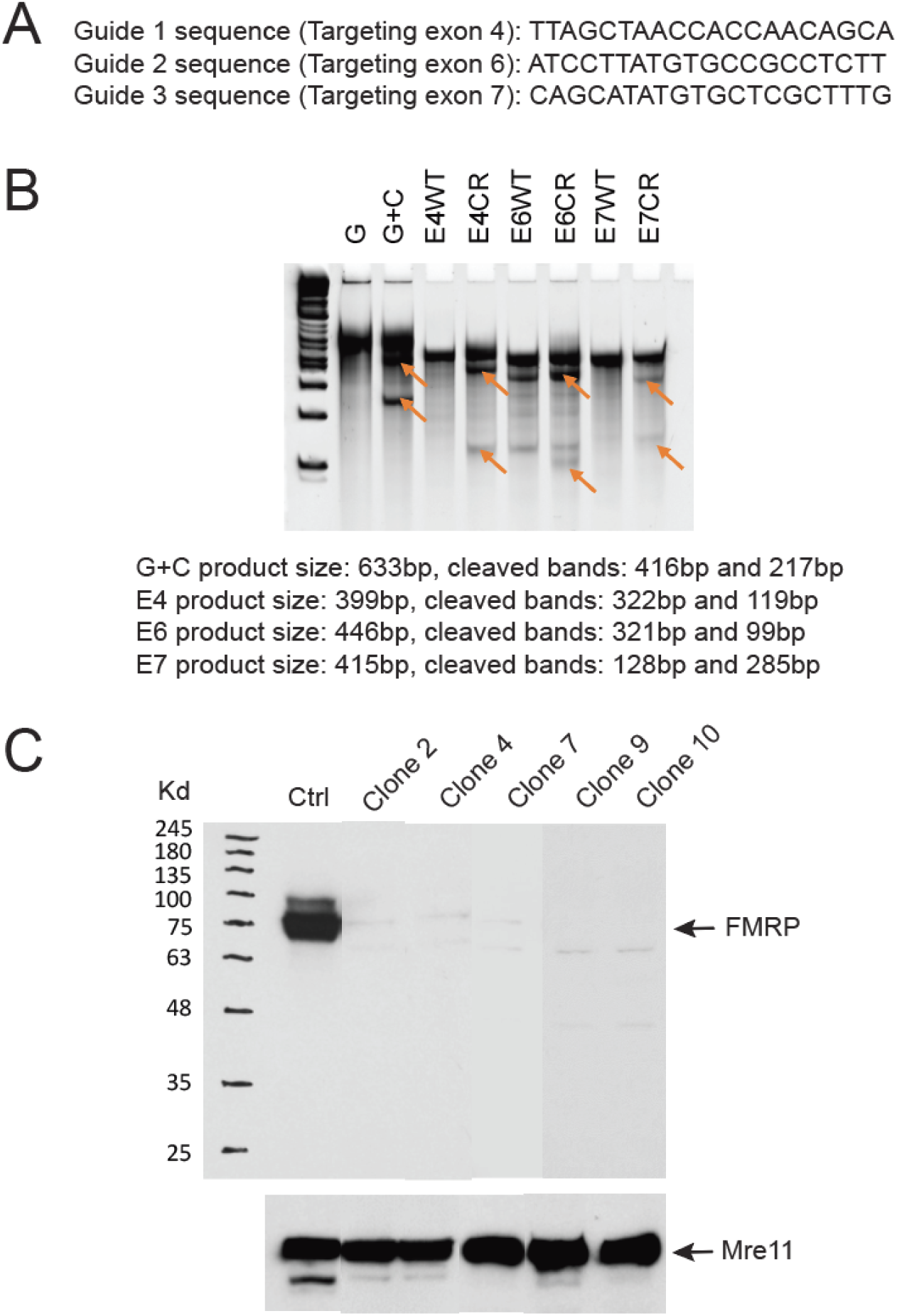
CRISPR knockout of *Fmr1* in tsA-201 cells. **A**, Three human *Fmr1* short guide RNA oligos targeting exons 4, 6 and 7 were designed, synthesized, ligated into the pX458 vector and confirmed by DNA sequencing. **B**, Electrophoresis result of the PCR products of genomic DNA fragments around exon 4, 6, and 7 from wild type cells (WT) and transfected cells (CR). The G+C DNA fragment was also amplified as positive control. All PCR products were cleaved by SURVEYOR Nuclease S following manufacturer’s protocol before being resolved in agarose gels. Cleaved bands were indicated by the arrows. **C**, Western blot of CRISPR knockout clones probed with a mouse anti-FMRP antibody (NBP2-01770, Novus Biologicals) and a rabbit anti- MRE11 antibody (NB100-142, Novus Biologicals) for control. Western blot result showed that *Fmr1* was knocked out in clones 2, 4, 7, 9 and 10.

**Figure 1-3.**
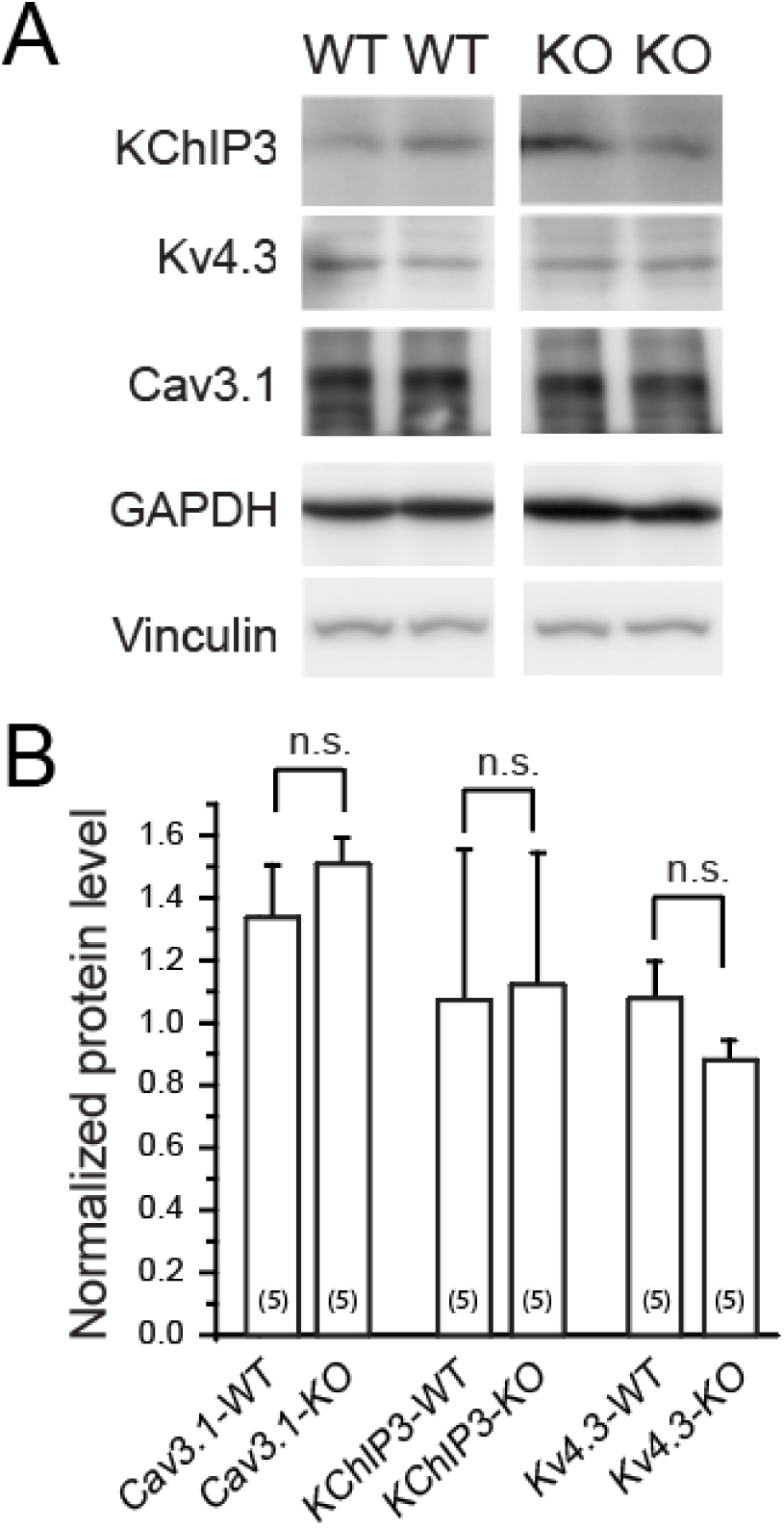
CRISPR knockout of *Fmr1* does not alter expression levels of subunits of the Cav3.1-Kv4.3- KChIP3 complex. Western blots and mean bar plots comparing the levels of Cav3.1-Kv4.3-KChIP3 subunits coexpressed in *Fmr1* WT and *Fmr1* KO tsA-201 cells. **A, B,** Representative Western blots (two replicates) (**A**) and mean bar plots (**B**) establish that the levels of each of the subunits are not significantly different between *Fmr1* WT and KO cells. In (**B**) the intensity levels of KChIP3 and Kv4.3 are normalized to the housekeeping gene GAPDH and Cav3.1 is normalized to Vinculin. Average values are mean ± S.E.M. with sample sizes (n) indicated in bar plots. n.s., not significant, Student’s two sample *t*-test.

**Figure 2-1.**
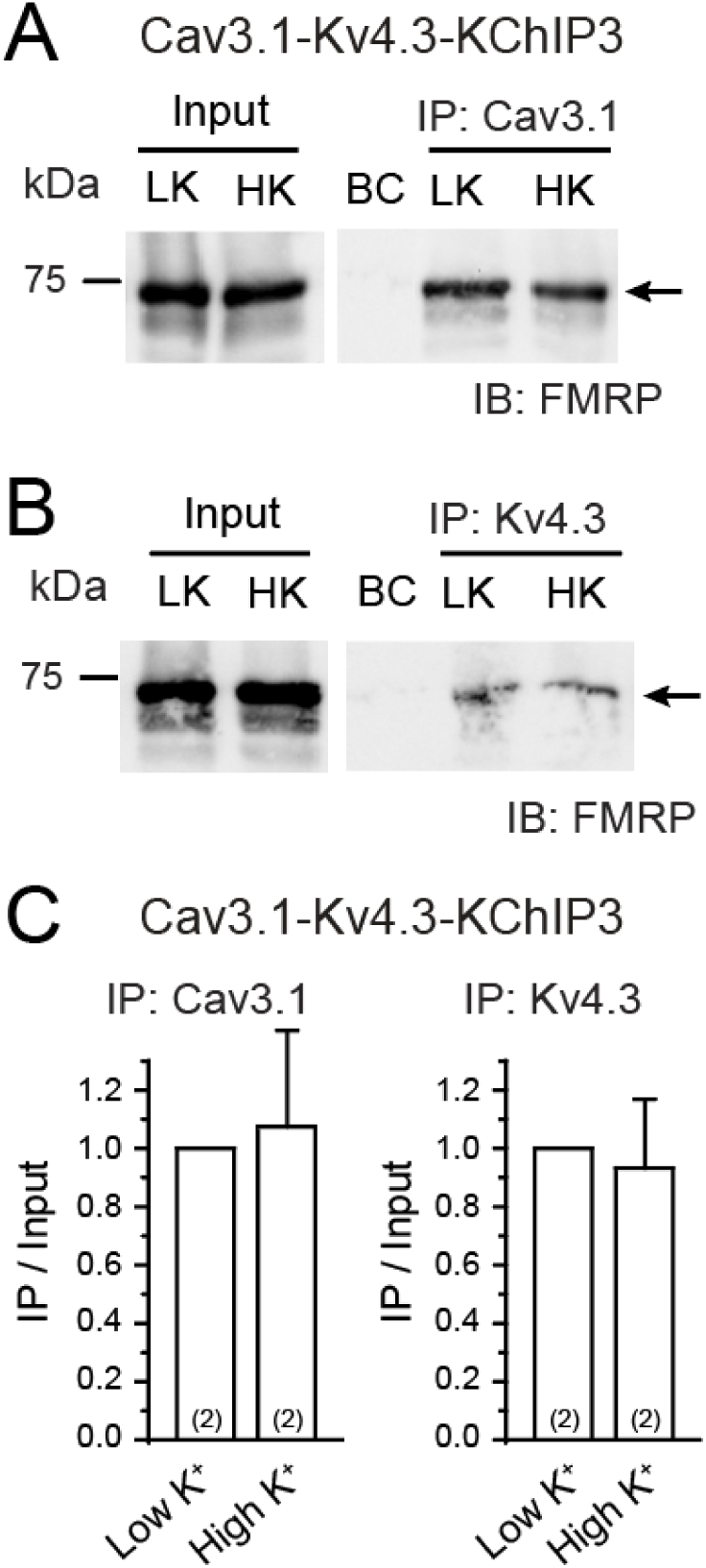
Cav3.1 calcium influx does not affect the co-IP between FMRP and either Cav3.1 or Kv4.3. A, B, Western blots of co-IP between Cav3.1/Kv4.3 and FMRP in WT tsA- 201 cells at a resting state of low [K]o (1.5 mM; LK) and 5 min after depolarizing cells with high [K]o (50 mM; HK). **C,** Mean bar graphs for the quantification of co-IP between FMRP and Cav3.1 (left) and Kv4.3 (right), as measured by the immunoprecipitated FMRP normalized to the level of FMRP in lysate. In contrast to the FMRP-KChIP3 relationship (Fig. 2E), membrane depolarization and calcium influx does not affect the co-IP between FMRP and either Cav3.1 (**A, C**) or Kv4.3 (**B, C**). Average values are mean ± SD with sample sizes (n) indicated in bar plots.

